# When is sympatric speciation a possible evolutionary outcome?

**DOI:** 10.1101/2023.05.31.543051

**Authors:** Pavithra Venkataraman, Supreet Saini

## Abstract

The process of speciation is the source of biodiversity. The most popularly accepted mode of speciation is allopatric speciation, where geography imposes the initial barrier to gene flow, and then biological barriers come up. On the other hand, sympatric speciation, which was not accepted as a possibility for long, requires that the process of speciation happen in the absence of a geographical barrier, in a well-mixed population. Several attempts have been made to theoretically identify the conditions in which speciation can occur in sympatry, but have several problems associated with them. We propose a model for sympatric speciation based on adaptation for resource utilization. We use this genetics-based model to investigate the relative roles of prezygotic and postzygotic barriers, from the context of ecological disruptive selection, sexual selection, and genetic architecture, in causing and maintaining sympatric speciation. We show that sexual selection that acts on secondary sexual traits does not play any role in the process of speciation in sympatry, and that assortative mating based on an ecologically relevant trait forces the population to show an adaptive response. We also demonstrate that understanding the genetic architecture of the trait under ecological selection is very important, and that it is not required for the strength of ecological disruptive selection to be very high in order for speciation to occur in sympatry. With this, we provide an insight into the kind of scenarios in which sympatric speciation can be demonstrated in lab.

## Introduction

Speciation is an important evolutionary outcome and understanding the phenomenology and genetics of speciation is a longstanding problem in biology. Historically, allopatric speciation, which is driven by geographical isolation is considered the null model of speciation[1–3]. The likelihood of the occurrence of sympatric speciation, where geography does not pose any hindrance to gene flow, was often thought to be an impossible phenomenon in nature [2, 3]. But, several empirical evidences of sympatric speciation in nature have been found [4–21], and thereafter, identifying the theoretical conditions and adjudging the likelihood of sympatric speciation has become important in the context of understanding the forces that drive generation of biodiversity.

In allopatric speciation, barrier to free exchange of genes in a population is first imposed by geography. Adaptive speciation theories suggest that the barrier to gene exchange due to biology comes up later, due to adaptive diversification [22]. But, in the case of sympatric speciation, geography plays no role in dictating the population’s evolutionary trajectory. Reproductive isolation has to occur “within a single interbreeding unit” [23] for sympatric speciation to occur. So, when is sympatric speciation a possible evolutionary outcome?

### Theories of sympatric speciation

Speciation, allopatric or sympatric, is the “fission of a gene pool” [24]. To explain the cause of this “fission” and its maintenance, several models have been proposed in the past few decades (reviewed systematically in [24, 25]). These models are based on some form of disruptive selection that acts on a trait of the individuals in the population. This idea was first proposed by Maynard Smith [26]. Because of the selection pressure imposed by disruptive selection, an adaptive response of the population is to diverge into two groups. But, in the absence of any geographical barrier, members of these two diverging groups are free to mate with each other, and produce unfit individuals. In order to prevent such “futile” mating events, a premating isolation said to evolve. This premating isolation in the form of preferential mating due to sexual selection has been studied extensively [27–34]. The preference of mates can be based on the trait under ecological selection, or selectively neutral marker traits. This preferential mating is called assortative mating. The genetic basis of assortativeness has been described both by one-allele [35–38] and two-allele models [35, 39].

Like sexual selection, disruptive selection, which is considered a basic driving force of speciation in sympatry, can be invoked to act on mating traits, secondary sexual characters, or physical traits that are ecologically relevant [26, 40–53]. If disruptive selection is due to an individual’s ecological interactions, due to limited availability of resources, it often operates in a frequency-dependent manner [22]. Typically, the population that inhabits a niche has a physical trait that determines its ability to utilize the available resources, and hence, its fitness. In such a population, this physical trait will evolve so as to maximize fitness in the given niche. In such a setting, frequency-dependent competition becomes an important determinant of an individual’s fitness [54]. This type of disruptive selection is called ecological disruptive selection.

While these broad frameworks to decipher the conditions in which sympatric speciation may occur exist, there are several problems associated with these models.

- Recent theoretical work shows that sympatric speciation can rarely ever occur only due to sexual selection [55].
- It has also been shown that the strength of disruptive selection required to drive and maintain speciation in sympatry is unnaturally high [56, 57].
- Several models depend on very strong assortative mating to drive divergence in the population, without any costs to the assortatively mating females [58].
- Assortative mating based on secondary sexual characters or neutral marker traits which have little ecological significance are used as traits on which disruptive sexual selection acts. The ecological relevance of such a selection pressure is an open question.
- The nature of genetic architecture of the traits that are under selection are extremely important. But, very few studies explore how variations in genetic architecture changes the likelihood of speciation in sympatry.

Based on these limitations, we develop a multi-locus genetic model to investigate the likelihood of sympatric speciation in a bird population. We invoke ecological disruptive selection on a physical trait (beak size), and sexual selection on female mating preference based on a male ornament, whose intensity depends on the male’s fitness. We investigate the relative roles of these two selection forces, and also ascertain how varying the genetic architecture and basis for assortative mating changes the evolutionary trajectory of the population. Our results show that assortative mating driven by sexual selection acting on secondary sexual characters does not play a role in generating genetic divergence. When the loci controlling the trait under ecological disruptive selection contributes unequally to the phenotype, we show that sympatric speciation can occur even when the strength of disruptive selection is low.

## Methods

### Model description

Imagine a bird population, where the fitness of an individual depends on its *beak size, x*. The environment that the birds are in, has two food resources (A corresponds to niche 1 and B to niche 2) that the birds can feed on. The variation of fitness with beak size is as shown in Figure 1A.

**Figure 1.**
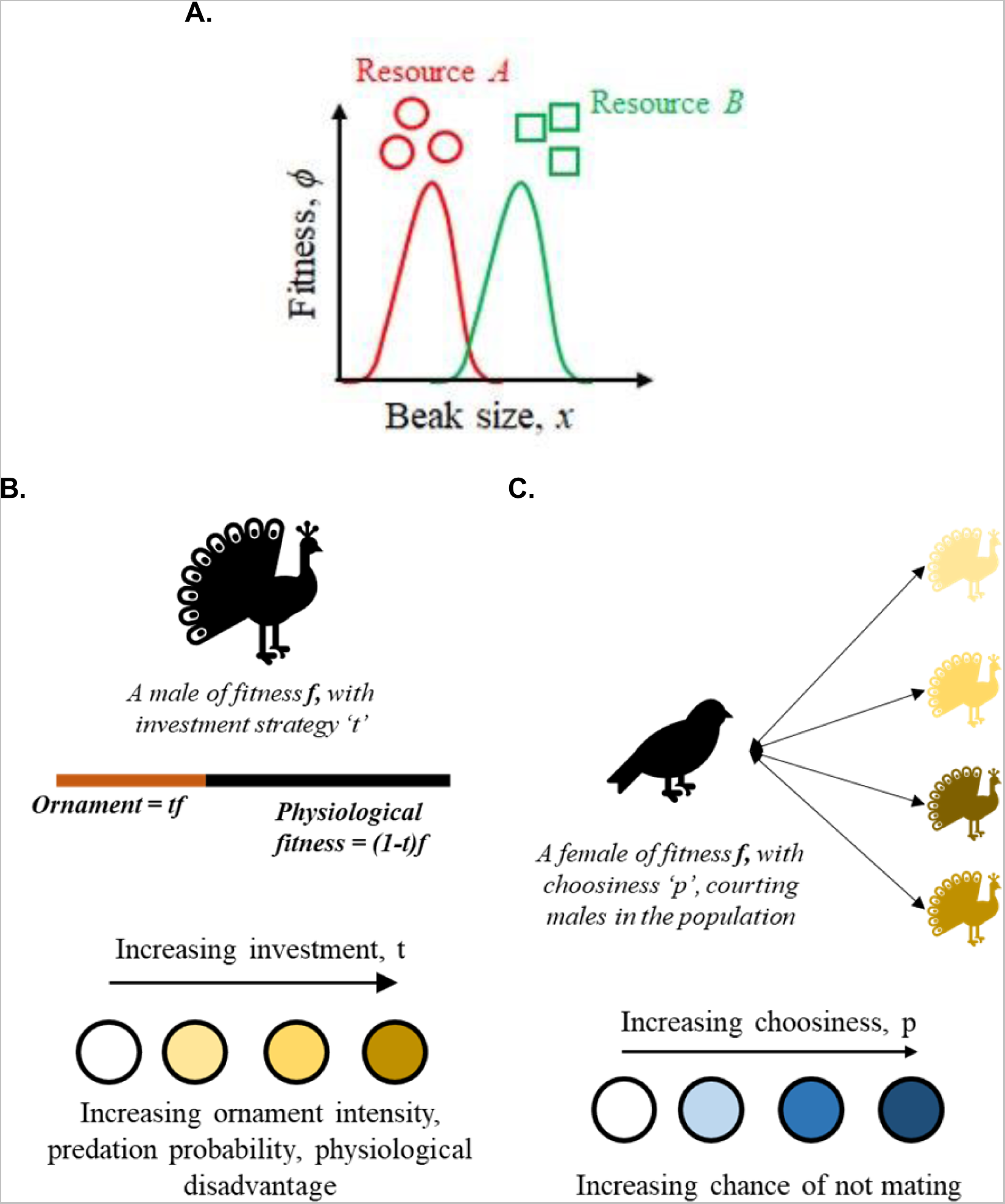
Model description. (A) The variation of fitness with beak size. In the hypothetical environment under consideration, there are two types of food resources, A and B. In order to utilize resource A, the beak size has to be small, while in order to effectively utilize resource B, the beak size has to be large. **(B-C) Behavior of the males and females.** In the bird population under consideration, the males invest a portion of their fitness to make an ornament, to attract the females. Increasing investment in making this ornament comes with costs including exposure to predators and a physiological disadvantage. Females, on the other hand, are choosy. They first court the males in the population and then pick a mate based on his ornament intensity. A highly choosy female may not find a partner.

The strength of ecological disruptive selection in this environment is quantified by the relationship in Equation 1.

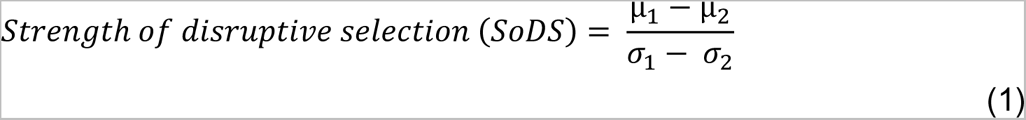

where ***μ****_i_* and ***σ_i_*** are the mean and standard deviation of the fitness distribution in ***i^th^*** niche.

Males in the environment are capable of investing a portion of their fitness in making an ornament to attract the females. The intensity of this ornament is given by Equation 2.

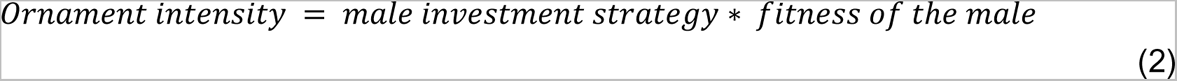

However, the males also incur costs for producing this ornament by increasing their exposure to predators, and by reducing their physiological fitness, as depicted in Figure 1B. Females, on the other hand, are capable of evolving “choosiness”, by virtue of which they preferentially pick their mates based on their ornament intensity. The ornament intensity is used by the females as a proxy of the fitness of the male it courts. A highly choosy female can accurately pick a male who showcases a high intensity ornament. But, such a female ends up with the risk of not finding a partner, as illustrated in Figure 1C. The females in the environment meet males in the environment, and the likelihood of this meeting is dependent on the physiological fitness of the males.

### Modeling Assortative Mating driven by Sexual Selection

Apart from ecological disruptive selection, the role of sexual selection in causing a prezygotic barrier via premating isolation mechanisms in the population is considered to be very important [59–61]. In this environment, we invoke sexual selection that is defined as an evolutionary force which influences mating success based on expression of certain traits [62]. It acts such that it drives assortative mating by enforcing a preference rule, based on a mating signal and a female’s preference for an elaborate mating signal. Such preference-based mating rules are common in theories that explain sexual selection [63–65].

Therefore, in this modeling setting, we invoke sexual selection to act on two traits – ornament intensity of the males, and the choosiness of the females. This is a scenario of assortative mating based on marker traits, which are not linked to the trait under ecological disruptive selection [66]. Both these traits are controlled by multiple loci.

### Modeling Assortative Mating based on Matching Rules

A mechanism that causes mating between likes, and hence leads to a prezygotic barrier, is called assortative mating. Typically, it is governed by preference rules (as explained above), or matching rules. Matching rules apply when the choosy individuals in a population (females in this case) inspect their own phenotype, and accordingly pick a male to mate with [67]. Such matching rules have been used in models that describe speciation [42, 68], and observed in ecology [69, 70]. The mate choice of the female is determined only by its own trait value, and is not governed by another trait (like choosiness; however, the strength of assortative mating can vary [67].

In our attempt to model assortative mating based on matching traits, we start from a non- assortatively mating population and check if its females evolve a bias towards mating with males whose beak size similar to theirs (and hence occupy the same niche). We call this bias the “assortativeness” of the females. This evolvable trait is controlled by ***n_a_*** number of unlinked equally contributing loci. Each locus contributes of ***δ_a_*** towards the female’s assortativeness. The mating bias of a female whose assortativeness is ***a*** is as explained in Equation 3.

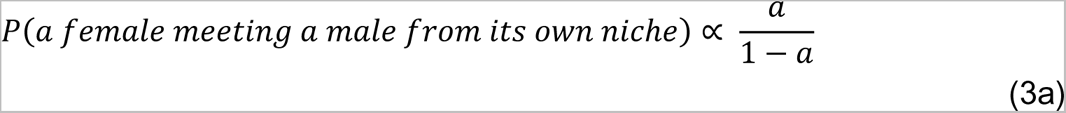

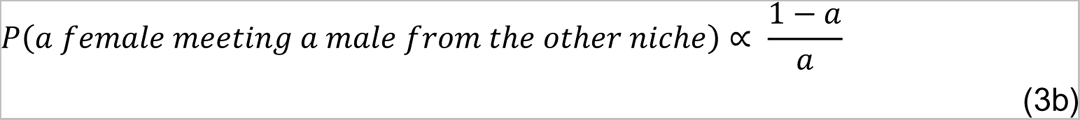

Thus, a highly assortative female runs into the risk of not finding a mate.

### Modeling Genetic Architecture of the Evolvable Traits

#### Beak size

The starting isogenic population which has a uniform beak size, ***x***, of 1. It is ill adapted to utilize either of the two resources because of ecological disruptive selection at play in the environment, and has to evolve beak sizes either smaller or greater than 1 to improve its fitness. The individuals in this population are diploids, that reproduce sexually. Beak size is a genetically determined phenotype, and is controlled by ***n_x_*** number of unlinked loci. Each locus contributes of ***δ_x_*** towards the beak size.

#### Male investment

There is sexual selection at play in the population, where the males exhibit an ornament to attract the females. In order to make the ornament, the males spend a fraction of their fitness. This fraction is called the *investment strategy***, *t***, and is a genetically determined phenotype. A total of ***n_t_*** number of loci control the investment strategy. Each of these ***n_t_*** number of unlinked loci contributes of ***δ_t_*** towards the investment strategy.

#### Female choosiness

A female’s mate choice is biased, based on the intensity of the ornament exhibited by a male. The greater the intensity, the greater the likelihood that the male finds a partner. This is because the females in the population are choosy. This *choosiness*, ***p***, is a genetically determined phenotype that is controlled by ***n_p_*** number of unlinked loci. Each of these ***n_p_*** loci contributes ***δ_p_*** towards determining the female’s choosiness.

All the loci in the population have only one of two alleles, ‘0’ or ‘1’. The value of a phenotype (***x*, *p*, *t***) is as shown in Equation 4.

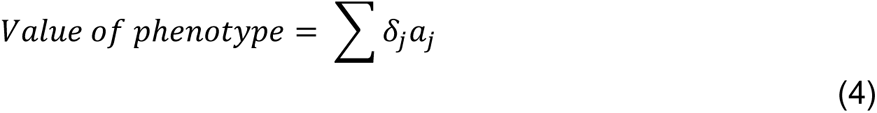

where *δ_j_* is the contribution of each allele towards a phenotype and ***a_j_*** is the numerical value (0 or 1) of the allele present at ***j^th^*** locus.

Therefore, the lowest value of a phenotype is 0 and the highest is ***2*n_loci_*δ***, where ***n_loci_*** is the number of loci controlling the phenotype, ***δ*** is the contribution of each locus to the phenotype.

Investing in making an ornament comes with certain costs to the male. First, the fitness available for growth and reproduction decreases as an investment towards making the ornament increases. If ***t*** is the fraction of fitness a male with fitness ***f_m_*** invests in making the ornament, the fitness available for physiological activities is shown in Equation 5.

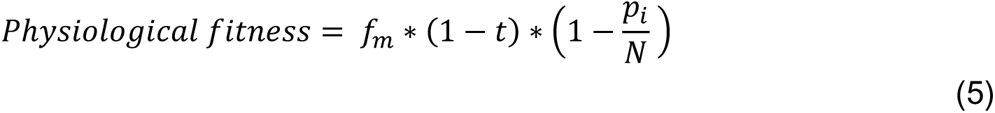

Where, ***p_i_*** is the number of individuals in the niche that the male occupies and **N** is the population size.

Second, the probability that the male is predated upon increases as the ornament intensity increases. The probability ***p_e_*_s_** is given by Equation 6.

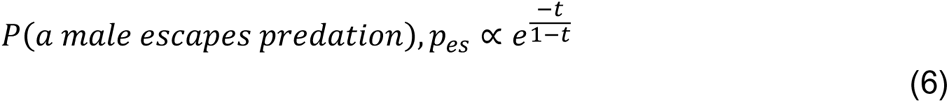

where ***t*** is the investment strategy.

The females in the population also pay a price for being choosy. As choosiness increases, the chance that a female finds a partner, decreases. This means that they do not get to pass on their genes to the next generation. The probability that a female finds a partner is given by Equation 7.

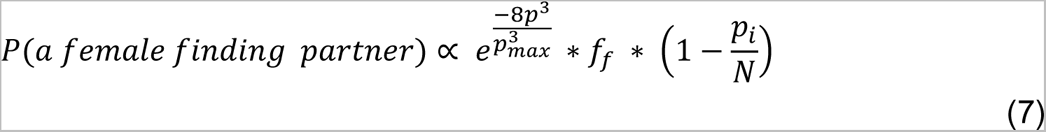

where, ***p*** is the choosiness, ***p_max_*** is the maximum choosiness genetically feasible and ***f_f_*** is the fitness of the female, ***p_i_*** is the number of individuals in the niche that the female occupies and **N** is the population size.

In this setting, the ability of a bird to utilize a food resource also depends on the competition it faces. The environment is such that if all the birds evolve either small or large beaks (and utilize only one of the food sources), their fitnesses decrease due to increased competition. The value of fitness of a bird of beak size ***x*** in the niche ***i*** is given by Equation 8.

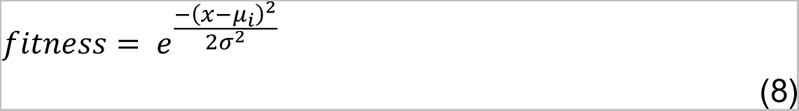

where ***μ_i_*** is the beak size of the fittest individual in niche ***i***, ***σ*** is the standard deviation of the distribution of fitness with beak size. ***n_i_*** (i=1,2) is the number of birds in niche ***i***, and **N** is the total number of birds in the population.

A female “meets” all the males in the population with a probability. The value of the probability is as shown in Equation 9.

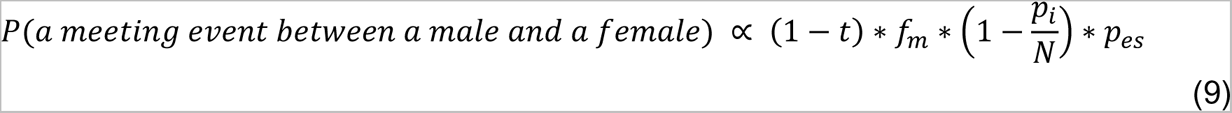

where ***t*** is the investment strategy of a male, ***f_m_*** is his fitness, and ***p_es_*** is the probability with which he escapes predation.

One of these “meeting” events leads to a “mating” event, the probability of which is explained by equation 10.

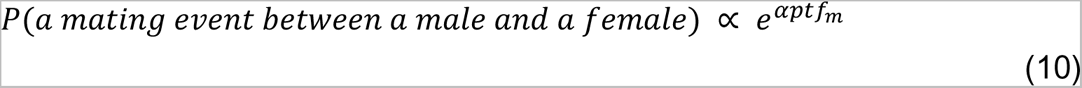

where, ***α*** is the strength of sexual selection in the environment, ***p*** is the choosiness of the female and ***tf_m_*** is the intensity of the ornament shown by the male.

One mating event leads to the production of one bird. Gender is assigned randomly. The generations are non-overlapping.

### Modeling of alterations in genetic architecture

#### a. Dominance

In the no dominance case, we model the contributions of the loci controlling a phenotype according to the following rules:

- Each of the two alleles at a given locus can either be ‘0’ or ‘1’.
- The contribution of each allele to the phenotype, when there are ***n_loci_*** number of unlinked loci governing it, is given by Equation 11.

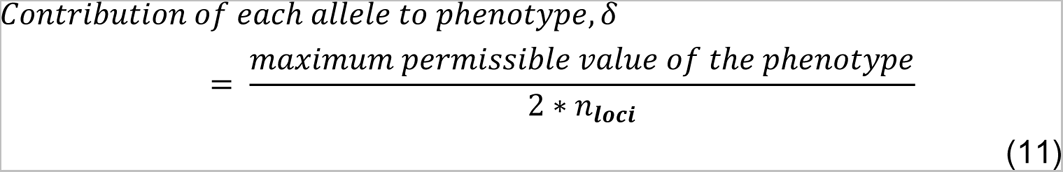

- The value of the phenotype is calculated as shown in equation 4.

In the dominance case

- Allele ‘1’ is considered to be dominant over allele ‘0’.
- So, we consider the contribution of a locus and not an allele (because one allele masks the presence of another), and is calculated as shown in Equation 12.

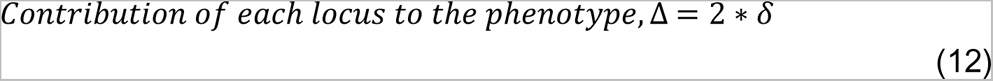

The contribution is modeled in this fashion to ensure that the maximum permissible beak size remains 2.

- If the number of loci with at least one ‘1’ is ***γ***, value of the phenotype is calculated as:

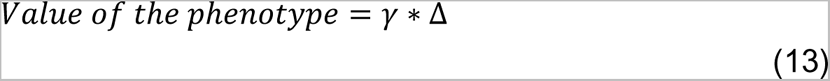

#### b. Unequal contribution

Quantitative traits are controlled by several loci that contribute small unequal values to the trait, and a few loci that contribute majorly to the value of the trait [71–73]. We model unequal contribution of the loci that control the three of the four evolvable traits – choosiness of the females, investment strategy of the males, and the beak sizes of individuals in the population- by drawing random numbers from an exponential distribution of a set mean, and normalize these values such that the minimum and maximum beak sizes genetically accessible are 0 and 2, respectively. The mean of the exponential distributions is equal to the contribution of each of the loci if they were contributing equally.

#### c. Unequal contribution and dominance

We also model a case where the loci that control a trait show dominance relationships and contribute unequally to the quantitative trait. In the case, we draw the values of contribution of each locus from an exponential distribution whose mean is twice the value of contribution of these loci when they were contributing equally to the phenotype, without showing dominance relationships. These values are then normalized to ensure that the minimum and maximum beak sizes genetically accessible are 0 and 2, respectively.

### Statistics

The error bars in the plots shown correspond to standard deviation, unless specified otherwise. Comparison of two means was done using a two-tailed t-test. A one-tailed t-test was used to ascertain if one mean is greater than or lesser than another, and the value of *d* used in these tests was 0.001. In all cases, the significance level was 0.05.

*p* values greater than 0.05, less than 0.05, less than 0.01, and less than 0.001 are indicated by **ns**, **\***, **\*\***, and **\*\*\*** respectively.

### Assumptions

● Only two alleles exist in the population at a given locus.
● Both alleles at a given locus stand a 50% chance of being passed on to the next generation.
● Mutations between these two alleles occur at a fixed rate.
● Loci controlling a trait are unlinked.
● No linkage exists between the loci controlling different traits.

### Simulation Steps

1. A female is chosen at random from the population, based on her choosiness and fitness.
2. The chosen female “meets” all the males in the population. The meeting probability is dependent only on the male’s fitness, as shown in equation 9.
3. A uniform random number is generated, and is used to pick one meeting event using Gillespie algorithm. The mating probability of the selected pair is calculated as given in equation 10. Another uniform random number is generated to decide if the mating event is successful or not, using Gillespie algorithm. Should a mating event take place, an offspring is born, and the population size increases by 1. The above steps are repeated until a fixed population size is reached. The first half of the individuals generated are females, while the second half are males.
4. Based on the beak sizes of individuals chosen for mating, the niche they occupy is decided. In a given niche, the frequency of different beak sizes is obtained and normal distribution is fitted with this data. Split in the population is quantified as shown in Equation 14.

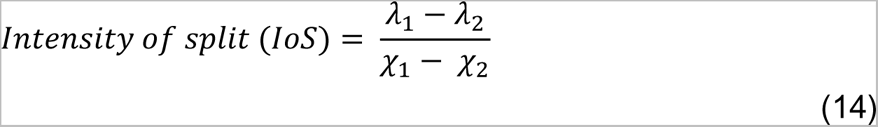

where ***λ_i_*** and ***χ_i_*** are the mean and standard deviation of the frequency distribution in the ***i^th^*** niche.

The intensity of split in the population is 0 even if there is only one individual of beak size 1 that is chosen for mating. Runaway selection is also a possibility in this modeling setting, especially if the strength of selection forces at play is very high. Since there is no coexistence of two types of individuals when runaway selection occurs, the value of intensity of split is considered to be 0.

In this work, the choosiness of the females and the investment strategy of the males is capped at 0.4 and 0.8 respectively.

All results, unless specified otherwise, are based on the data obtained after simulations for 50 generations. All simulations were performed in MATLAB R2019a. The codes used in the study are provided in the Supplement.

## Results

### An isogenic population adapts to split into two phenotypically heterogeneous groups

We start with a population where each individual has the same genotype for beak size, and hence the same beak size (=1). Similarly, every female has the same genotype for choosiness, and every male’s genotype for investment strategy is identical. The starting population’s females are not choosy (choosiness = 0), and the males do not make the ornament (investment strategy = 0). Each of the loci that control *x, p,* and *t* contribute equally to the phenotype. The strength of disruptive selection is 3.84, and the strength of sexual selection is 5 This will be considered as the “null case” henceforth, unless specified otherwise.

In this modeling setting, there are two types of adaptive responses that are possible – (a) the entire population shifting to one of the niches, which we call runaway selection, and (b) the population splitting into two (equally or unequally) and occupying both the niches, which we term as sympatric speciation. We consider a population to be ill-adapted even if one of the individuals chosen for mating (among the 2000 individuals that mate) has a beak size equal to 1.

The frequency distribution of the beak size in the population at the start and end of 50 generations are shown in the Figure 2A and Figure 2B respectively. The frequency distribution of female’s choosiness and male’s investment strategy at the end of 50 generations are shown in Figure 2C and Figure 2D respectively.

**Figure 2.**
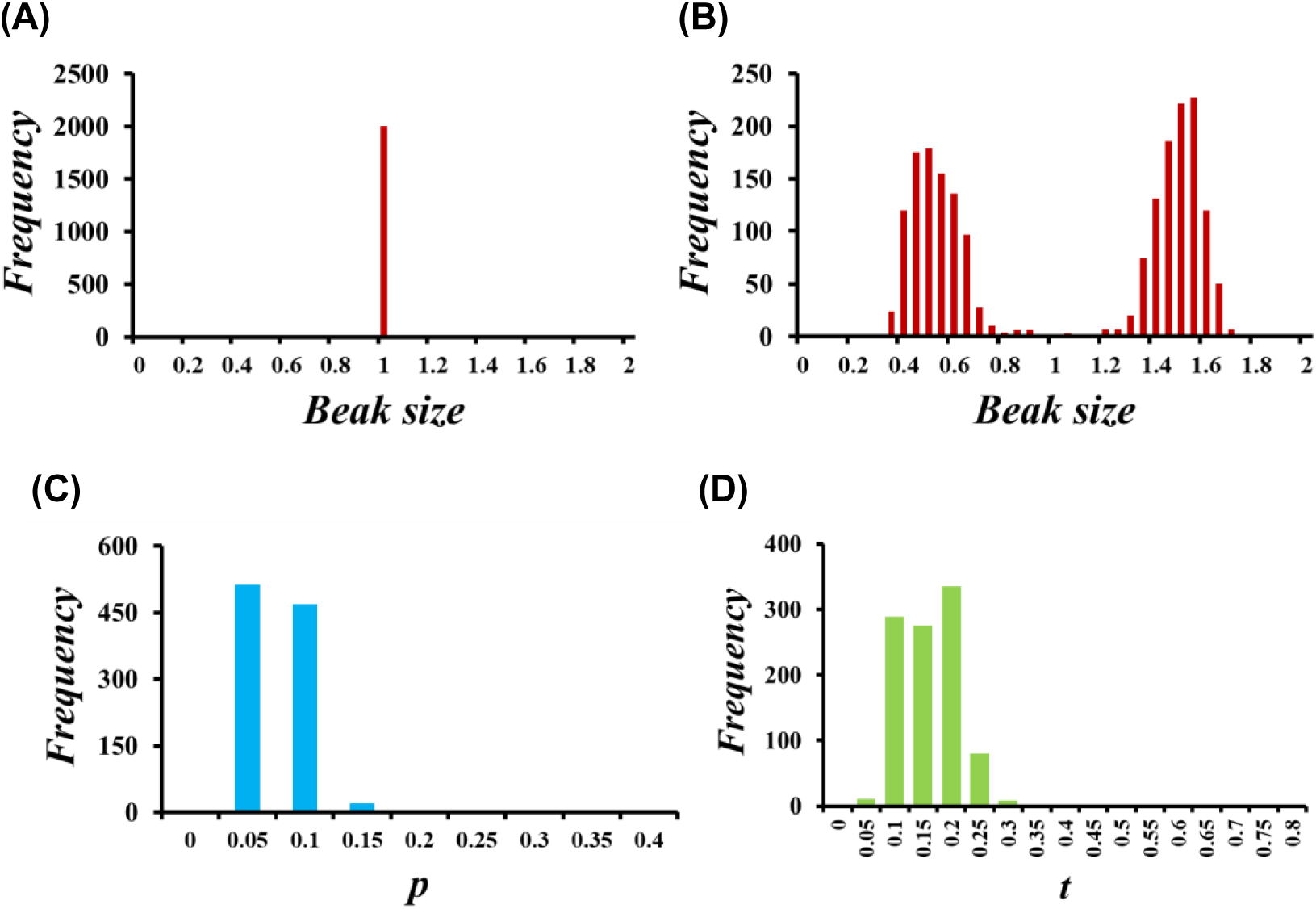
An isogenic population splits into two groups in 50 generations. In our modeling setting, we start with a population of 1000 individuals that has a uniform beak size, as shown in **(A)**. When the strength of ecological disruptive selection is ∼3.84 and the strength of sexual selection is 5, we see that the due to post-zygotic barriers, the population splits into two groups at the end of 50 generations, as shown in **(B)**. The intensity of this split is calculated to be ∼5.1157. **(C)** and **(D)** shows the frequency distribution of the choosiness of the females, and the investment strategy of the males in the population, respectively.

### Increasing the strength of disruptive selection increases the likelihood and intensity of population split, along with the possibility of runaway selection

In this modeling setting, individuals whose beak sizes are 0, 1, and 2 have the lowest fitness. Other beak sizes confer a fitness to the birds depending on the strength of disruptive selection. When disruptive selection is low, it is possible for the individuals in the population to acquire a significant gain in fitness without making a big change to their beak sizes. But, as the strength of disruptive selection increases, small deviations from the starting beak size do not increase fitness of the individual significantly. So, in order to survive in environments where the disruptive selection is strong, the individuals in the population could (a) evolve to have beak sizes less than 1, and more than 1, and speciate in sympatry, or (b) evolve to have beak sizes less than 1, or more than 1, and undergo runaway selection.

Our results show that as the strength of disruptive selection increases, the population’s tendency to undergo a split increases, up to a certain extent, as shown in Figure 3A, along with the split intensity (quantified in equation 14), as shown in Figure 3B. (For Figure 3B, only cases where the population underwent a split have been considered for calculation of the average split. So, for the case when strength of disruptive selection is ∼1.67, there does not exist a value of intensity of split.) But, with further increase in the strength of disruptive selection, there is increasing pressure on the population to show an adaptive response, and therefore, the likelihood of the population to undergo runaway selection increases. Therefore, the intensity of split at high values of strength of disruptive selection, when calculated considering all the possibilities, drops as depicted in the heat plots in Figure S1. The effect of changing the strength of disruptive selection, on beak size, choosiness of females, and investment strategy of the males, at the end of 50 generations, are shown as heat plots in Figures S2-S4.

**Figure 3.**
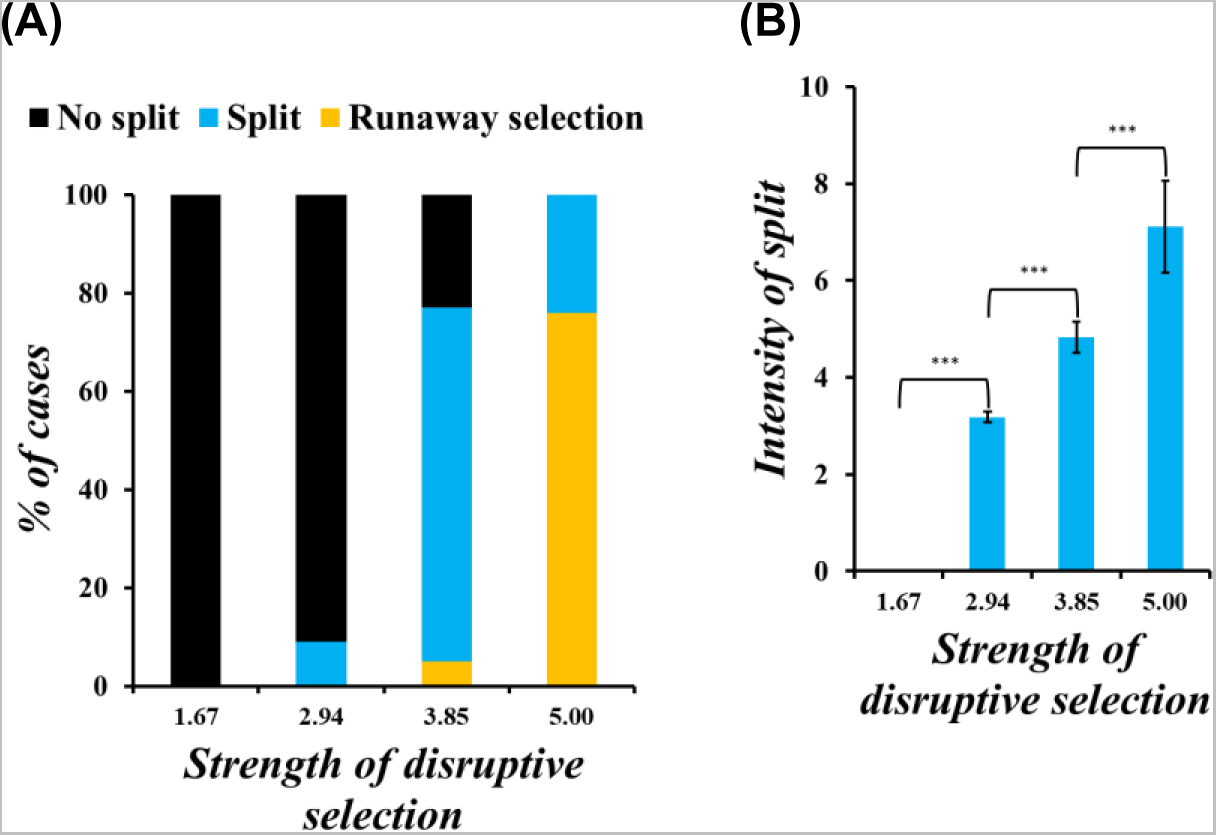
An increase in the strength of disruptive selection increases the tendency of the population to split, the intensity of split, and the likelihood of runaway selection. At low values of strength of disruptive selection, the population does not split into two. As the strength increases, the likelihood of the population to split increases, along with the intensity, as shown in **(A)** and **(B)**. But, as depicted in **(A)**, when the strength of disruptive selection increases, the population’s tendency to undergo runaway selection also increases. We check if (i) the IoS (SoDS=1.67) < IoS (SoDS=2.94), (ii) if IoS (SoDS = 2.94) < IoS (SoDS = 3.85), and (iii) if IoS (SoDS = 3.85) < IoS (SoDS = 5), using one-tailed t-tests. The p-values obtained for all three tests are less than 0.001. Since these p-values obtained indicate high significance, we conclude that IoS (SoDS =1.67) < IoS (SoDS = 2.94) < IoS (SoDS = 3.85) < IoS (SoDS = 5)

This non-monotonic behavior of the population as the strength of ecological disruptive selection increases can be explained by looking at the genetic architecture of the loci that control the beak size of the birds in the population. At low values of strengths of disruptive selection, even small deviations from that starting beak size (=1) are significantly beneficial. This means that the frequency distributions of the beaks centered around 0.5 and 1.5 have a large width. But, as the intensity of strength of disruptive selection increases, small deviations from the intermediate beak size are no longer significantly beneficial. In such a case, the population has to evolve beak sizes closer to either 0.5 or 1.5, thus reducing the width of the distribution of beaks centered around 0.5 and 1.5. What increasing the strength of disruptive selection also does, is that the population’s trajectory becomes increasingly dependent on the number of high fitness males and females occupying a given niche in the first few generations. This dependence leads to an increase in probability of runaway selection as the strength of disruptive selection increases. Population dynamics obtained after simulations, when the strength of disruptive selection is ∼2.94, ∼3.85, and 5 are shown in Figure 4, Figure 5, and Figure 6 respectively.

**Figure 4.**
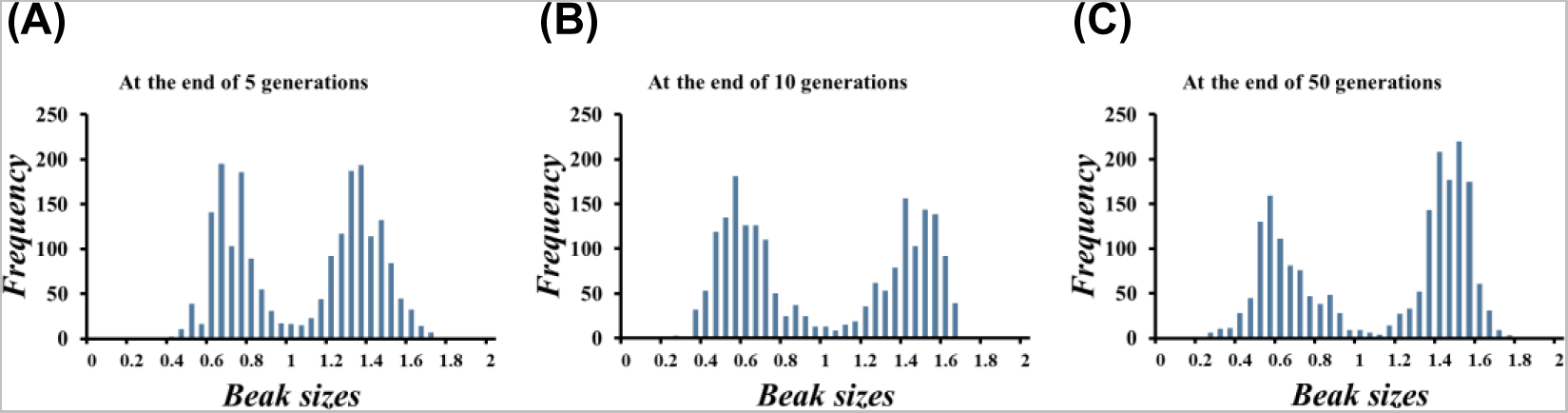
Population dynamics when the strength of disruptive selection is low. In these three figures, the distribution of beak sizes of the individuals of the population is shown at three time points – after 5 generation **(A)**, after 10 generations **(B)**, and after 50 generations **(C)**. Clearly, when the strength of ecological disruptive selection is low (∼2.94), the population does not split at the end of 50 generations.

**Figure 5.**
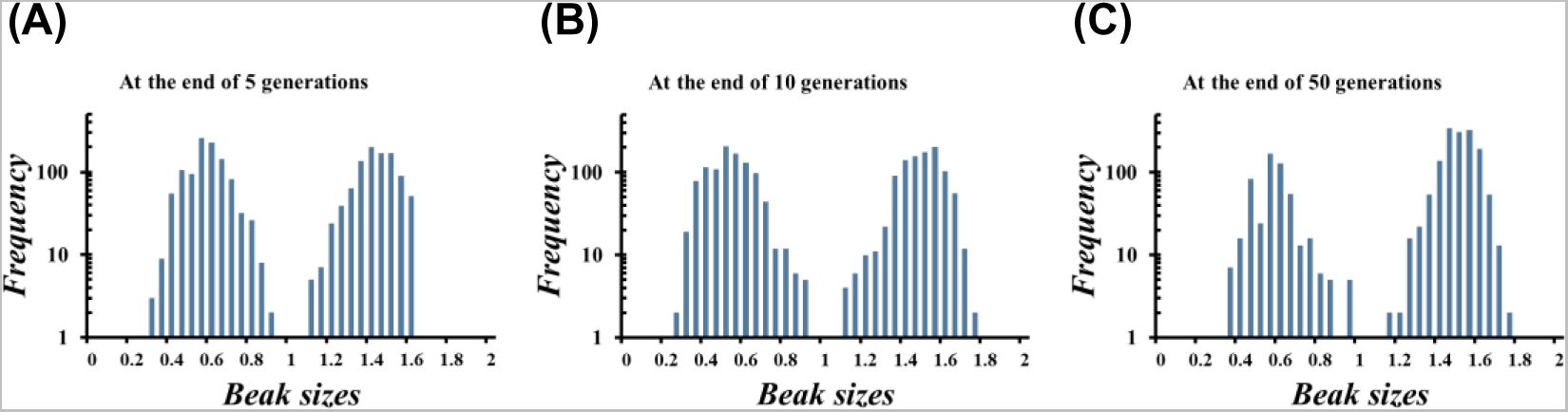
Population dynamics when the strength of disruptive selection is intermediate. In the three figures above, the frequency distribution of beak sizes on the individuals in the population is shown after 5 generations **(A)**, after 10 generations **(B)**, and after 50 generations **(C)**. At the end of 50 generations, the population has undergone a split when the strength of ecological disruptive selection is ∼3.85.

**Figure 6.**
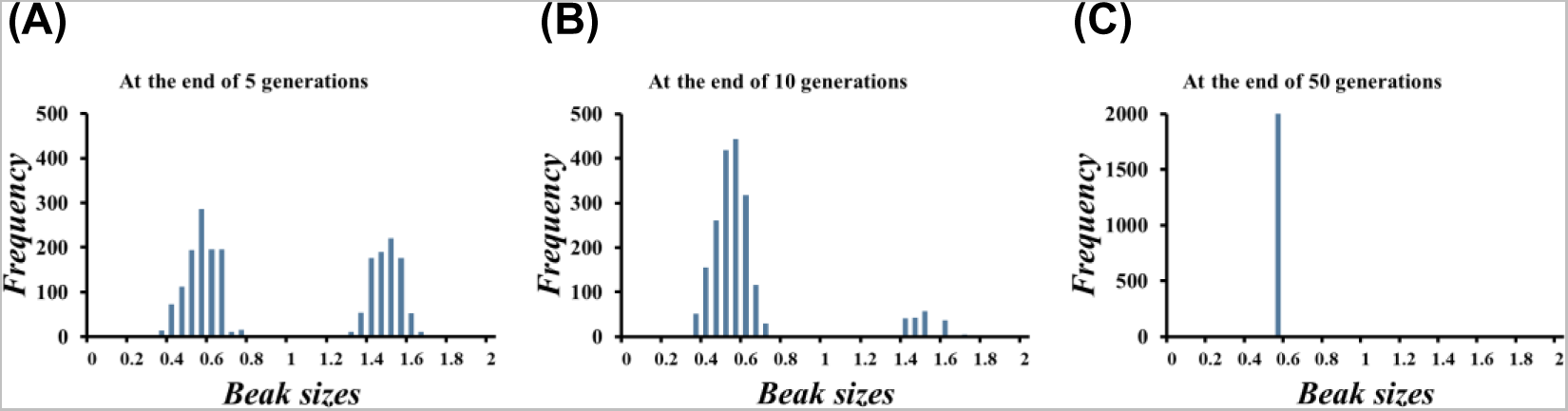
Population dynamics when the strength of disruptive selection is high. As described in the text, the evolutionary trajectory of a population experiencing strong disruptive selection would depend on the number of high fitness individuals occupying both the niches. In **(A)**, the number of individuals in both the niches is almost the same. But, as time progresses, the number of fit individuals occupying one niche is much higher than the other **(B)**, and eventually, the population shifts in that direction **(C)**. The strength of disruptive selection was 5.

### Strength of sexual selection does not have any role to play in dictating the population behavior

Several models describe how divergence of mating preferences drives speciation in sympatry [45, 74–77]. But, it is has been argued in the past that it is very rare for such a divergence to arise in an isogenic or identical population, and be maintained [75, 77, 78]. In a starting population like in our model, given that there is ecological disruptive selection acting on beak sizes, mating bias of females is more likely to be based on a trait that is reflective of a male’s fitness, than on a magic trait.

We consider sexual selection to be acting on two marker traits - ornament size of the males and the choosiness of the females. The males in population can evolve to produce an ornament to attract the females. But, making this ornament is costly - the males pay a fitness cost, and expose themselves to predators. The females, on the other hand, may not find a mate if they become too choosy. Since the type of ornament produced by males from both the niches is the same, there is no divergence in mating preferences shown by the females.

When sexual selection acts on these two traits in this setting, our results show that it does not play any role in causing speciation in sympatry, as shown in Figure 7A and Figure 7B. (These results are obtained when the strength of disruptive selection ∼3.85. See Figure S1 for data of split obtained in other cases). In fact, sexual selection may lead to a loss in genetic divergence by bringing together a choosy female from one niche and a male with an intense ornament from the other niche. The effect of changing the strength sexual selection, on beak size, choosiness of females, and investment strategy of the males, at the end of 50 generations, are shown as heat plots in Figures S2-S4.

**Figure 7.**
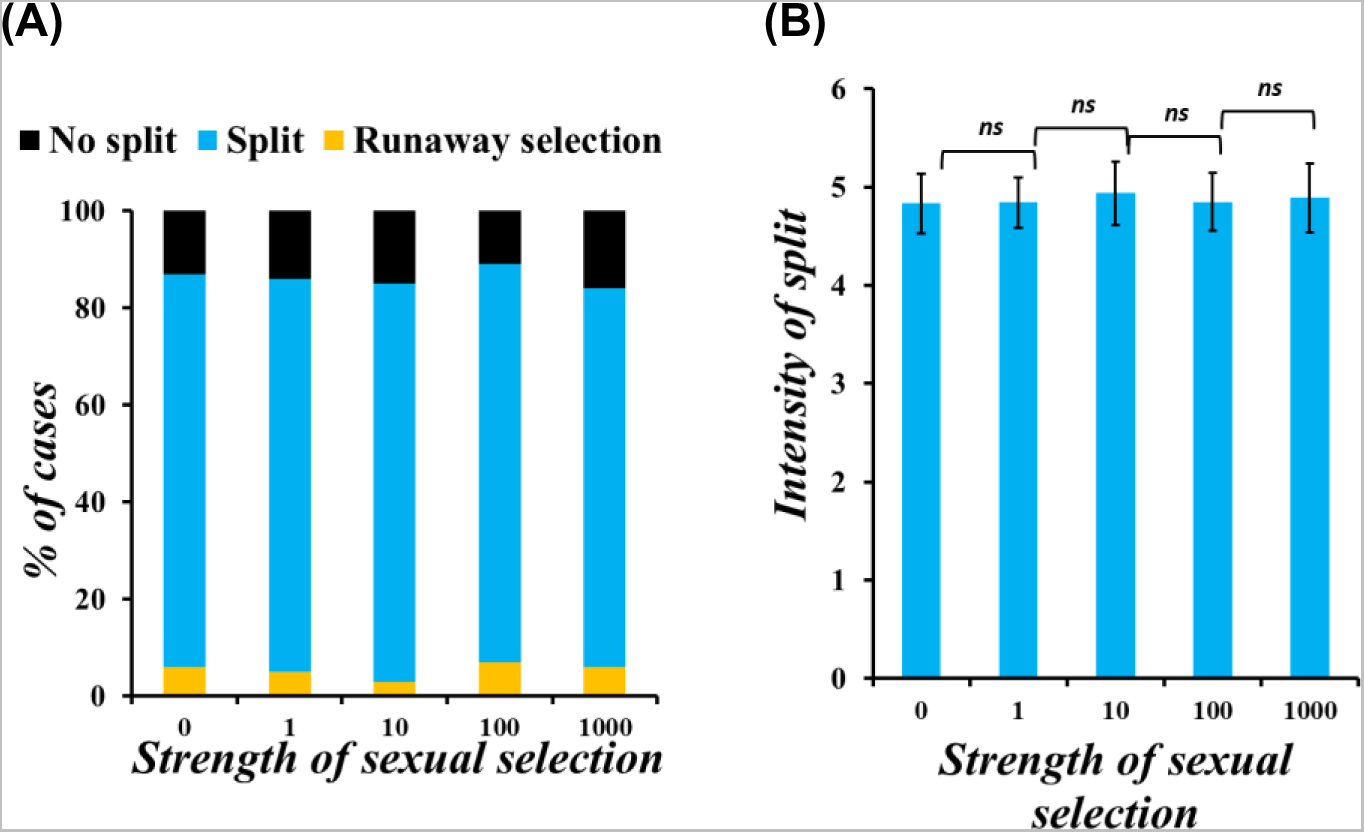
Sexual selection does not play a role in dictating the evolutionary trajectory of the population. In this modeling setting, sexual selection acts on two secondary sexual characters - choosiness of the females, and ornament size of the males. The figure describes the results obtained when the strength of disruptive selection is ∼3.85. **(A)** The proportion of cases where the population undergoes a split, runaway selection, or does not split remains the same with increase in the magnitude of strength of sexual selection at play. **(B)** The intensity of split, whenever it occurs, also does not change with change in the strength of sexual selection. The error bars indicate the standard deviation of the data.

We next investigate the effect on the split intensity when (a) the females are not choosy, (b) when the males do not invest in making an ornament, and (c) when the females are not choosy and the males do not invest in making an ornament. The results when the strength of disruptive selection is 3.85 and the strength of sexual selection is 5 are as shown in Figure 8.

**Figure 8.**
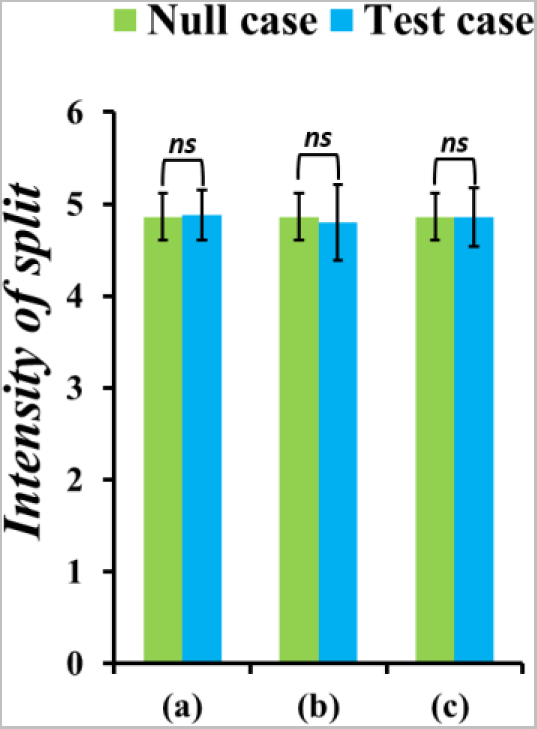
There is no effect on the intensity of split in the population when the secondary sexual traits of the females and males are absent. We check if the choosiness of the females or the investment strategy of the males has any role in dictating the evolutionary trajectory of the population. Null case here corresponds to the scenario where the females are choosy, and the males invest in making an ornament. **(a)** In an environment where the females are not choosy but the males invest in making an ornament, there is no change in the intensity of split in the population (*p=*. **(b)** When the males do not invest in making an ornament but the females are choosy, there is no change in the intensity of split in the population. **(c)** When the females are not choosy and the males do not make an ornament, the intensity of split in the population does not change.

When the females in the environment are not choosy, and/or the males in the environment do not invest in making an ornament, the intensity of the split in the population remains the same as that in the null case, where the females are choosy, and the males are making an ornament. This shows that sexual selection and the traits that it acts on have no role to play in the maintenance of genetic divergence in this setting.

### Assortative mating enforces an adaptive response

In order to understand the role of a pre-zygotic barrier in our model, we allow females in the population to evolve assortative mating based on beak size. This means that when the females mate assortatively, a female bird with a “small” beak will prefer to mate with a male with a “small” beak, and vice versa. The value of assortativeness can range between 0 and 1. We check the effect of introducing assortativeness at different values of strength of disruptive selection.

Our results show that the population either splits or undergoes runaway selection in the presence of assortatively mating females. While it was shown in Figure 2 that increase in strength of disruptive selection increases the population’s tendency to undergo runaway selection, this tendency increases significantly when the females mate assortatively. This is seen when one compares Figure 9A with Figure 9B.

**Figure 9.**
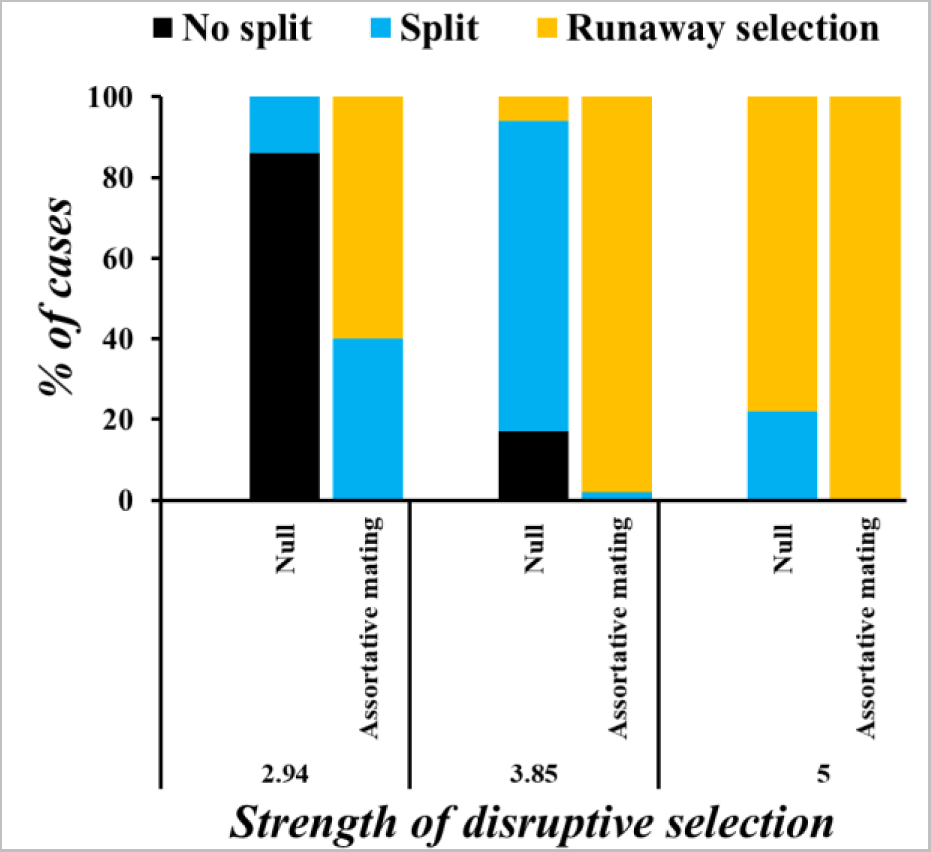
Assortatively mating females drive the population to undergo a split, or runaway selection. **(A)** shows the behavior of the population when there is no assortative mating, and **(B)** when there is assortative mating based on beak sizes in the population. It is seen that even when the strength of disruptive selection is low, there is an adaptive response shown by the population in the presence of assortative mating. This adaptive response is either runaway selection, or split. With increase in the strength of disruptive selection, the proportion of cases where runaway selection occurs is more in the presence of assortative mating than when it is absent.

### Assortative mating increases the intensity of split in the population

In the presence of assortatively mating females, we understand that the likelihood of runaway selection is high than when there is no assortative mating. But, when the population splits, it is seen that the mean split is higher when there is assortative mating exhibited by the females, as seen in Figure 10. For Figure 10, only cases where the population underwent a split have been considered for calculation of the average split. So, for the case when strength of disruptive selection is 5, there does not exist a value of intensity of split. For the other two cases, a one tailed t-test was performed to check if the increase in split intensity with increase in strength of disruptive selection is significant. The results indicate that the increase in split due to assortative mating is highly significant.

**Figure 10.**
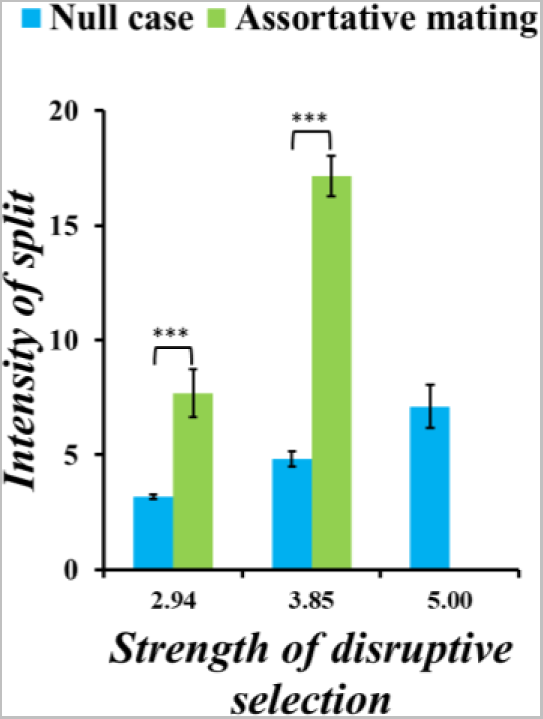
Assortative mating increases the intensity of split in the population. In the presence of assortatively mating females, whenever the population splits, it is seen that the intensity of split is higher than when the females don’t mate assortatively.

**Figure 11.**
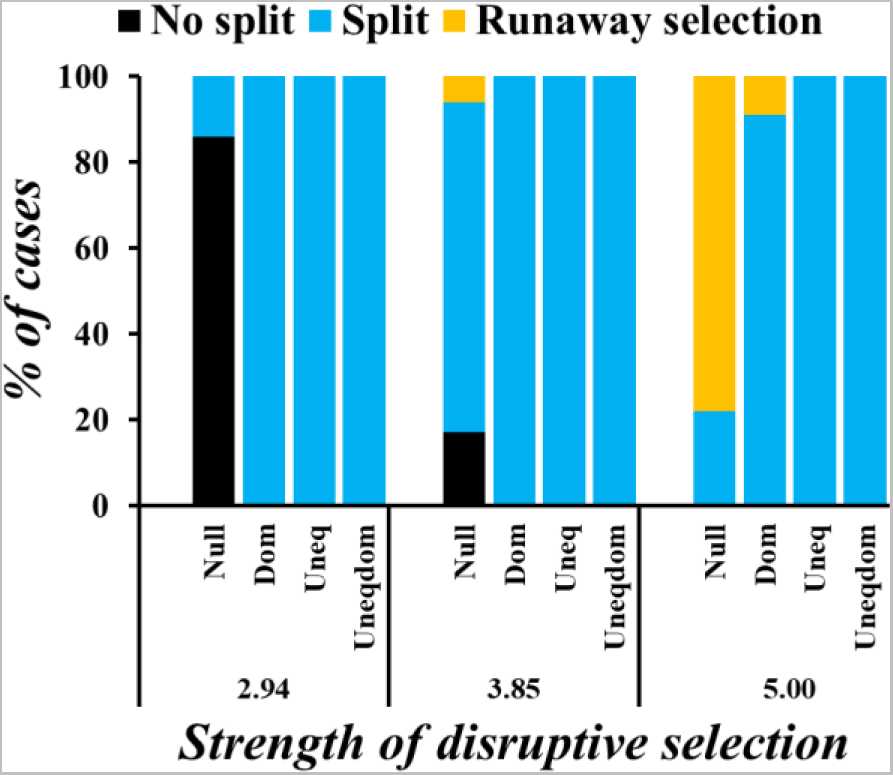
The likelihood of the population to undergo runaway selection decreases when there is a change in the genetic architecture. Results are shown for data obtained when the strength of sexual selection is 5. The graphs show the proportion of cases where the population does not split, undergoes runaway selection, or a split when **(A)** the loci controlling beak size do not show dominance and contribute equally to the trait, **(B)** when they show dominance relationships, **(C)** when they contribute unequally, and **(D)** when they contribute unequally and have dominance relationships governing them.

This increase in intensity of split is because, when individuals of the same niche mate repeatedly, genetically accessible beak sizes reduce in number. In the presence of disruptive selection, the beak sizes of the individuals of a given niche are very similar, thereby increasing the intensity of split.

Since the evolvable traits in the population are genetically controlled, we next sought to investigate the role of genetic architecture in determining the population’s evolutionary trajectory. Since the loci controlling the three evolvable traits are not linked, we vary their nature independently and check the population’s adaptive response.

### Introduction of dominance and unequal contribution individually and in combination in the loci controlling beak size reduces the likelihood of runaway selection

In the loci that control beak size, we make alterations to the genetic architecture by introducing (a) dominance, (b) unequal contributions, and (c) unequal contributions and dominance. Our results, as depicted in Figure 11, show that the likelihood of runaway selection decreases in all the three cases. The strength of sexual selection used to obtain data for Figure 11 is 5. Beak size and intensity of split do not change with increase in strength of sexual selection when the nature of loci controlling beak size is altered (Figures S5-S12).

In the case where are dominance relationships exist, the number of accessible beak sizes is less, but one step change in the genetic makeup confers a 2x advantage (in the no dominance case, an allele going from 0 to 1 increases the beak size by 0.05, but in the dominance case, it increases it by 0.1). This eases the possibility of the population to split, by creating fit individuals in both the niches, in lesser time. So, the phenomenon of the population moving in one direction reduces.

In the case where the loci contribute unequally with or without dominance relationships, the number of accessible beak sizes increases, when compared to the null case where the loci contribute equally and show no dominance (since contributions of the loci are drawn from an exponential distribution, beak sizes like 0.59989 may be possible, compared to when they had to be multiples of 0.05 in the null case). As a result, creation of fit individuals in both the niches is easier, and as a result, the likelihood of the population to split into two increases.

Both dominance relationships and unequally contributing loci enhance the possibility of acquiring beak sizes that enable survival in the presence of ecological disruptive selection.

### Conclusions and Discussion

Sympatric speciation is proposed as one of the possible ways using which species biodiversity on earth can be explained. But, what causes sympatric speciation, when does it occur, and what facilitates speciation in sympatry is not well understood. Several models have been able to demonstrate sympatric speciation in theory, by making a number of assumptions that often cannot be explained using first principles in biology [22, 24, 25, 36, 55–58, 79]. For instance, why would disruptive selection act strongly on traits which are not reflective of an organism’s phenotypic or genotypic quality is not very clear. Similarly, divergence in the mating preferences of the individuals (mostly females) in the population could only under very rare conditions in nature. In several models, assortative mating based on preferences is used to demonstrate sympatric speciation. However, these preferences come with costs, that are often not accounted for. Also, the relative roles of the forces that cause speciation in sympatry, and genetic architecture of traits under selection are not very clearly understood.

Using our model, we show speciation in sympatry and explain the contributing forces and their roles. In our model, there is ecological disruptive selection acting on the beak sizes of the individuals of a bird population. These birds are diploid and reproduce sexually. We invoke sexual selection to act on two secondary sexual traits - ornament size of the males, and choosiness of the females. The ornament intensity of the male is modeled such that it reflects the individual’s genotype quality. Since the starting population is isogenic for all the evolvable traits, we propose that it is unlikely that a divergence in mating preference based on secondary sexual traits, in order to survive the ecological disruptive selection will arise, at least until the population has undergone some genetic divergence in beak size [80]. In the absence of such a divergence in mating preference that constitutes a pre-zygotic barrier, we demonstrate that a split in the population can arise and be maintained with the help of post-zygotic barriers that exist due to the ecological selection pressure.

We then investigate the role of strength of disruptive selection in dictating the evolutionary outcome of the population. Our results show that as strength of disruptive selection acting on the population increases, its tendency to show one of the two adaptive responses – split, or runaway selection – increases. An increase from low values of strength of ecological disruptive selection increases the tendency of the population to split, but with more increase, the population is more likely to undergo runaway selection. This is in spite of accounting for competition within a niche. But, the tendency to undergo runaway selection in one direction is seen only when the loci controlling the beak size of the birds contribute equally to the phenotype. When they show dominance relationships and/or contribute unequally, we see that this tendency decreases drastically, and is almost lost in most cases as shown in Figure 8.

A common critique associated with invoking disruptive selection to demonstrate sympatric speciation is that it is often required to be very strong (and that is not very common in nature) in order to maintain the split in the population. But, we have been able to show that even when the strength of disruptive selection is low, split in the population can occur and be maintained when the loci controlling the trait under selection do not contribute equally to the phenotype. It must be noted that several complex traits (like beak size) are said to be controlled by many loci whose contributions are small, and a few large-contribution loci [81]. So, we believe that the proposition that genotypic divergence can occur when ecological disruptive selection is acting on a continuous quantitative trait controlled by quantitative trait loci is not unreasonable. This result not only provides an insight into the type of experiments that can be designed to demonstrate sympatric speciation, but also shows the importance of understanding the genetic architecture of the loci that control the trait under disruptive selection.

As discussed before, there are problems associated with modeling sexual selection based on traits that are not an indicator of an organism’s quality of genotype [82–84]. We address this issue by invoking sexual selection to act on a trait – an ornament - whose intensity is dependent on the organism’s fitness. We also account for the costs that come with producing such an ornament. Similarly, the females in the population are allowed to evolve choosiness based on this ornament. Sexual selection acts on these two traits such that it helps in producing a fit offspring, by increasing the probability of mating between fit males and females. We report that in the absence of a divergence in the marker traits produced by the males, sexual selection seems to have no role in dictating the population’s evolutionary fate. This result is in contradiction to the one reported in the past [42] that assortative mating based on neutral marker traits can lead to a split in the population, and does not agree with a result reported previously [80] that divergence in mating preferences is not required for the population to speciate in sympatry. In fact, in this modeling setting, sexual selection in the environment tends to homogenize the genetic diversity in the population by favoring the mating of a very choosy female and a male with a bright ornament from different niches. A hybrid produced by this mating faces post-zygotic barriers because of its low fitness and does not survive. The following strategies can prevent these “futile” cross-niche matings - (a) the individuals evolve niche specific ornaments, and (b) the individuals mate assortatively based on the trait that is under ecological disruptive selection (beak size in this case). Exhibiting ornaments come with two costs for the male - increased risk of predation, and lower fitness left to perform physiological activities. On the other hand, evolving assortativeness based on the beak size comes with the cost of not being able to find a mate. Although this population is under frequency-dependent selection, stabilizing selection could act to push all individuals to one of the two optimum beak sizes, especially when the strength of ecological disruptive selection is very high. Therefore, assortativeness as a strategy to impose a pre-zygotic barrier is more likely to evolve in this population. We tested this by invoking a strong mating bias in the females, and saw that the tendency of the population to undergo runaway selection increases. While the modeling of bias can be done in several ways, it is important to note the significance of this result - the number of fit assortatively mating females occupying a given niche in the initial generations dictates the population’s trajectory.

The nature of pre-zygotic barriers that will arise in this population to prevent cross-niche matings remains to be clearly ascertained. Also, we are yet to determine how this strategy that poses a pre-zygotic barrier will evolve to get linked to the trait under ecological disruptive selection.

In our model, the females in the population can mate as many times as they want, and we understand that is not representative of events that occur in nature. But, we consider a population of 1000 individuals in our simulations, and believe that constraining the number of mating events of a female may only quantitatively change the population’s behavior, and not qualitatively. We also start with a population that is isogenic for the trait under ecological disruptive selection. There are two simple ways by which the starting population can be made non-isogenic – (a) there is a continuous distribution of individuals between beak size 0 and 2, and (b) there is a normal distribution centered around 1. In the first case, we believe that genetic divergence is going to be very likely, and the likelihood of runaway selection is almost zero, since there are equal number of fit individuals occupying both the niches. However, in the second case, the population’s fate is most likely to depend on the number of fit males and females that occupy both the niches in the first few generations, and the population will move to the niche where there are more fit individuals.

Although the choosiness and assortativeness of the females, and the investment strategy of the males of the starting population is zero, the entire population is not isogenic for these two traits (females and males differ in the loci that control choosiness, assortativeness and investment strategy). This quickens the process of evolution of these traits. If we were to start with a population that was isogenic for all these traits, and if evolution were to occur only via mutations, the population would take much longer to attain the values of these phenotypes that we report.

We are yet to ascertain the effects of introducing assortative mating based on matching rules when the loci controlling the trait under disruptive selection do not contribute equally. The effect of varying the number of loci controlling these traits is also not very clear as yet.

Although sympatric speciation has been thought of an important route via which species diversity can arise, it has not been demonstrated experimentally. Our results provide an insight into an experimental design that can be adopted to demonstrate speciation in sympatry.

## Supporting information

Simulation Code

## Acknowledgements.

This work was funded by a DBT/Wellcome Trust (India Alliance) grant (Award No. IA/S/19/2/504632) to SS. PV is supported by the Prime Minister’s Research Fellowship (PMRF ID 1302050).

