## Supplementary material for "When is sympatric speciation a possible evolutionary outcome?": Simulation Code

### Codes

mainfile

```
clc
close all

dim1=1;
dim2=20; %number of loci controlling x,t, and p
xmu=2/(2*dim2); %contribution of each locus controlling x
tmu=0.8/(2*dim2); %contribution of each locus controlling t
pmu=0.4/(2*dim2); %contribution of each locus controlling p
totp=2*dim2*pmu; %maximum permissible choosiness

%unequal contributions of loci controlling x
uneqx1=exprnd(xmu,dim1,dim2);
uneqx=2*uneqx1/sum(uneqx1);

%dominance + unequal contribution of loci controlling x
uneqxdom11=exprnd(2*xmu,dim1,dim2/2);
uneqxdom1=uneqxdom11/sum(uneqxdom11);
uneqxdom12=exprnd(2*xmu,dim1,dim2/2);
uneqxdom2=uneqxdom12/sum(uneqxdom12);
uneqxdom=[uneqxdom1, uneqxdom2];

%unequal contribution of loci controlling p
uneqp1=exprnd(pmu,dim1,dim2);
uneqp=0.4*uneqp1/sum(uneqp1);

%dominance + unequal contribution of loci controlling x
uneqpdom11=exprnd(2*pmu,dim1,dim2/2);
uneqpdom1=uneqpdom11/sum(uneqpdom11);
uneqpdom12=exprnd(2*pmu,dim1,dim2/2);
uneqpdom2=uneqpdom12/sum(uneqpdom12);
uneqpdom=0.2*[uneqpdom1, uneqpdom2];

%unequal contribution of loci controlling t
uneqt1=exprnd(tmu,dim1,dim2);
uneqt=0.8*uneqt1/sum(uneqt1);

%dominance + unequal contribution of loci controlling t
uneqtdom11=exprnd(2*tmu,dim1,dim2/2);
uneqtdom1=uneqtdom11/sum(uneqtdom11);
uneqtdom12=exprnd(2*tmu,dim1,dim2/2);
uneqtdom2=uneqtdom12/sum(uneqtdom12);
uneqtdom=0.4*[uneqtdom1, uneqtdom2];

%computation of values of intensity of split
spec=zeros(1,1,16);
dev_split=zeros(1,1,16);
beaks=zeros(1,1,16);
choose=zeros(1,1,16);
invest=zeros(1,1,16);
dev_beaks=zeros(1,1,16);
dev_choose=zeros(1,1,16);
dev_invest=zeros(1,1,16);
```

```
page=1;
for i=1:4
    for j=1:4
        for k=1:4
            if i==1
               bi=2;
               dellx=xmu;
       else if i==2
            bi=1;
            dellx=2*xmu;
       else if i==3
                bi=2;
                dellx=uneqx;
       else if i==4
                 bi=1;
                 dellx=uneqxdom;
                end
            end
        end
            end
    if j==1
        pi=2;
        dellp=pmu;
        sump=0.4;
    else if j==2
            pi=1;
            dellp=2*pmu;
            sump=0.4;
        else if j==3
                pi=2;
                dellp=uneqp;
                sump=sum(uneqp);
            else if j==4
                    pi=1;
                    dellp=uneqpdom;
                    sump=sum(uneqpdom);
                end
            end
        end
    end
         if k==1
        ti=2;
        dellt=tmu;
    else if k==2
            ti=1;
            dellt=2*tmu;
        else if k==3
                ti=2;
                dellt=uneqt;
            else if k==4
                    ti=1;
                    dellt=uneqtdom;
                end
            end
```

```
            end
             end

[spec1,splitstdev,bavg,pavg,tavg,devb,devp,devt]=qfic(dim1,dim2,dell
x,dellp,dellt,sump,bi,pi,ti);
        spec(:,:,page)=spec1;
        dev_split(:,:,page)=splitstdev;
        beaks(:,:,page)=bavg;
        choose(:,:,page)=pavg;
        invest(:,:,page)=tavg;
        dev_beaks(:,:,page)=devb;
        dev_choose(:,:,page)=devp;
        dev_invest(:,:,page)=devt;
        page=page+1;
        end

    end
end


function[split,stdevsplit,avgbeak,avgp,avgt,stdev_beak,stdev_p,stdev
_t]=qfic(sz1,sz2,xdell,pdell,tdell,psumm,ib,ip,it)

%width of distribution of resources
stdev1=0.13;
stdev2=0.13;

n1=length(stdev1);

%strength of sexual selection
stsex=5;
n2=length(stsex);

split=zeros(n1,n2);
sep=zeros(n1,n2);
itersep=zeros(n1,n2,50);
beaks=zeros(2000,50);
choosiness=zeros(1000,50);
investments=zeros(1000,50);

divsel=zeros(n1,1);

for i=1:n1
    devst1=stdev1(i);
    devst2=stdev2(i);
    divsel(i)=1/(devst1+devst2); %strength of disruptive selection
     for j=1:n2
        alp=stsex(j); %strength of sexual selection
        iter=1;
        while iter<=50
        [sep(i,j),finalbeaks,pees,tees]=
symsp(devst1,devst2,alp,xdell,pdell,tdell,psumm,sz2,ib,ip,it);
        spval(iter)=sep(i,j);
        devt(iter)=std(tees);
        meant(iter)=mean(tees);
```

```
        for m=1:n1
            for n=1:n2
                if sep(m,n)<0
                    sep(m,n)=0;
                end
            end
        end

        split(i,j)=split(i,j)+sep(i,j);
        itersep(i,j,iter)=sep(i,j);
        beaks(:,iter)=finalbeaks;
        choosiness(:,iter)=pees;
        investments(:,iter)=tees;
        iter=iter+1;
        end

avgbeak(i,j)=mean(mean(beaks)');
bavg=avgbeak(i,j);
avgp(i,j)=mean(mean(choosiness)');
pavg=avgp(i,j);
avgt(i,j)=mean(mean(investments)');
tavg=avgt(i,j);

sumb=0;
sumt=0;
sump=0;

for m=1:2000
    for n=1:50
        sumb=sumb+((beaks(m,n)-bavg)^2);
    end
end

for m=1:1000
    for n=1:50
        sumt=sumt+((investments(m,n)-tavg)^2);
        sump=sump+((choosiness(m,n)-pavg)^2);
    end
end

stdev_beak(i,j)=sqrt(sumb/(2000*50));
stdev_t(i,j)=sqrt(sumt/(1000*50));
stdev_p(i,j)=sqrt(sump/(1000*50));
     end
end

stdevsplit=std(itersep,0,3);
split=split/50;
end

function[diff,matxf,choosi,invest]=symsp(sig1,sig2,alpha,delx,delp,d
elt,sump,nloci,bi,pi,ti)

nind=1000;  %number of individuals
ngenerations=50; %number of generations
```

```
matm=zeros(nind,1); %array of males that mate%
matf=zeros(nind,1); %array of females that mate%

xtrack=zeros(ngenerations,nind); %beak size of individuals of every
generation%
ttrack=zeros(ngenerations,nind/2); %investment strategies of the
males of every generation%
ptrack=zeros(ngenerations,nind/2); %choosiness of the females of
every generation%
mtrack=zeros(ngenerations,nind); %investment strategy of males of
every generation
ftrack=zeros(ngenerations,nind); %choosiness of females of every
generation

fpcr1=zeros(nind/2,nloci); %first strand of female's gene - p
fpcr2=zeros(nind/2,nloci); %second strand of female's gene - p
mpcr1=zeros(nind/2,nloci); %first strand of male's gene - p
mpcr2=ones(nind/2,nloci); %second strand of male's gene - p

ftcr1=zeros(nind/2,nloci); %first strand of female's gene - t
ftcr2=ones(nind/2,nloci); %second strand of female's gene - t
mtcr1=zeros(nind/2,nloci); %first strand of male's gene - t
mtcr2=zeros(nind/2,nloci); %second strand of male's gene - t

if bi==2
fxcr1=zeros(nind/2,nloci); %first strand of female's gene - x
fxcr2=ones(nind/2,nloci); %second strand of female's gene - x

mxcr1=zeros(nind/2,nloci); %first strand of male's gene - x
mxcr2=ones(nind/2,nloci); %second strand of male's gene - x


else if bi==1
fxcr1=zeros(nind/2,nloci); %first strand of female's gene - x
fxcr2=zeros(nind/2,nloci); %second strand of female's gene - x
mxcr1=zeros(nind/2,nloci); %first strand of male's gene - x
mxcr2=zeros(nind/2,nloci); %second strand of male's gene - x
%altered genetic makeup in case of dominance
fxcr2(1:nind/2,1:nloci/2)=1;
mxcr2(1:nind/2,((nloci/2)+1):nloci)=1;
end
end

%means of normal distributions centered that indicate disruptive
selection%
mean1=0.5;
mean2=1.5;

for i=1:ngenerations

    ngen(i)=i;


[ff,fm,xavg,xvar,xtot,np1,np2,fx,mx]=fcalc(fxcr1,fxcr2,nind,mxcr1,mx
cr2,nloci,delx,mean1,mean2,sig1,sig2,bi); %function that calculates
beak size and fitness of the individuals
```

```
    avgxx(i)=xavg;
    varxx(i)=xvar;
    fitness=cat(1,ff,fm);
    avgf(i)=mean(fitness);

    femfit=ff;
    malfit=fm;

    [fp]=pcalc(fpcr1,fpcr2,nind,nloci,delp,pi); %function that
calculates choosiness of every female%
    avgp(i)=mean(fp);

    %calculation of probability that a female finds a partner based
on its
    %fitness and choosiness
    fcost=(exp(-8*fp.*fp.*fp/(sump)^3)).*femfit';
    sumfcost=sum(fcost);
    fprob1=fcost/sumfcost;

    sumf=0;
    for x=1:nind/2
        sumf=sumf+fprob1(x);
        fprob(x)=sumf;
    end

    [mt]=tcalc(mtcr1,mtcr2,nind,nloci,delt,ti); %function that
calculates investment strategy of every male%
    avgt(i)=mean(mt);

    % calculation of probability that a male escapes predation%
    for jj=1:nind/2
        predprop(jj)=exp(-mt(jj)/(1-mt(jj)));
    end

    summcost=sum(predprop);
    predprob=predprop/summcost;
    summ=0;
    for y=1:nind/2
        summ=summ+predprob(y);
        mprob(y)=summ;
    end

    %assignment of beak sizes, males' investment strategy and
females'
    %choosiness
    if i>=1
        xtrack(i,:)=xtot;
        ttrack(i,:)=mt;
        ptrack(i,:)=fp;
    end

    %meeting and mating event determination
    for p=1:nind
        r1=rand;
        ii=1;
        while r1>fprob(ii)
```

```
            ii=ii+1;
        end
        matf(p)=ii;

        if i==ngenerations
        choosi(p)=fp(ii);
        end

        for q=1:nind/2
            fdep(q)=(1-(mt(q)))*malfit(q)*mprob(q); %product of the
selected female's fitness with all the males' fitness
            matingp(q)=exp(mt(q)*malfit(q)*fp(ii)*alpha); %mating
probability of the selected female with all the males in the
population
        end
            sumfdep=sum(fdep);
            fdepp=fdep/sumfdep;
            summatp=sum(matingp);
            probmating=matingp/summatp;

            sumffdep=0;
            summating=0;

            for pp=1:nind/2
              sumffdep=sumffdep+fdepp(pp);
              fitnessprob(pp)=sumffdep;
              summating=summating+probmating(pp);
              matprb(pp)=summating;
            end

        iiii=1;
        while iiii==1
            r2=rand;
            r3=rand;
            iii=1;
         while r2>fitnessprob(iii)
            iii=iii+1;
         end
         ev=iii;
         if r3>matprb(ev)
            iiii=1;
         else
             iiii=0;
         end
        end
        matm(p)=ev;
        if i==ngenerations
        invest(p)=mt(ev);
        end
    end

    for b=1:nind
        indf=matf(b);
        indm=matm(b);
        matfemx(b)=fx(indf);
        matmalx(b)=mx(indm);
```

```
    end

    if i>=1
        mtrack(i,:)=matmalx;
        ftrack(i,:)=matfemx;
    end



    %determination of the genotypes of the offspring produced by the
mating event%

[fempcr1,fempcr2,malpcr1,malpcr2]=pmat(fpcr1,fpcr2,mpcr1,mpcr2,nind,
nloci,matf,matm);

[femtcr1,femtcr2,maltcr1,maltcr2]=tmat(ftcr1,ftcr2,mtcr1,mtcr2,nind,
nloci,matf,matm);

[femxcr1,femxcr2,malxcr1,malxcr2]=xmat(fxcr1,fxcr2,mxcr1,mxcr2,nind,
nloci,matf,matm);

    %assignment of female and male genotypes of the next generation%
    fpcr1=fempcr1;
    fpcr2=fempcr2;
    mpcr1=malpcr1;
    mpcr2=malpcr2;

    ftcr1=femtcr1;
    ftcr2=femtcr2;
    mtcr1=maltcr1;
    mtcr2=maltcr2;

    fxcr1=femxcr1;
    fxcr2=femxcr2;
    mxcr1=malxcr1;
    mxcr2=malxcr2;
end

femxf=ftrack(ngenerations-1,:);
malxf=mtrack(ngenerations-1,:);
matxf=[femxf malxf];
femalep=ptrack(50,:);
malet=ttrack(50,:);

%calculation of intensity of split
diff=1;
bs1=0;
for pp=1:(2*nind)
    if matxf(pp)==1
        bs1=bs1+1;
    else
        bs1=bs1+0;
    end
end

if bs1~=0
```

```
    diff=0;
else
nsmallx=0;
nbigx=0;
 for hh=1:(nind*2)
    if matxf(hh)<1
        nsmallx=nsmallx+1;
    else if matxf(hh)>1
            nbigx=nbigx+1;
    end
    end
 end

  if nsmallx==0 && nbigx~=0
    diff=-1;
  else if nsmallx~=0 && nbigx==0
          diff=-1;
  else if nsmallx<2 || nbigx<2
          diff=-1;
  else if (nsmallx~=0 && nbigx~=0) && (nsmallx>2 && nbigx>2)

     smallx=zeros(nsmallx,1);
     bigx=zeros(nbigx,1);

     fmatx=sort(matxf);

     smallx=(fmatx(:,1:nsmallx))';
     bigx=(fmatx(:,(nsmallx+1):end))';

     pd1=fitdist(smallx,'Normal');
     m1=mean(pd1);
     dev1=std(pd1);
     pd2=fitdist(bigx,'Normal');
     m2=mean(pd2);
     dev2=std(pd2);

     diff=(m2-m1)/(dev2+dev1);
  end
  end
  end
  end
end

end


function[fitf,fitm,avgx,varx,totx,p1,p2,xf,xm]=fcalc(xfcr1,xfcr2,ind
n,xmcr1,xmcr2,loci,xdel,meano,meant,sigma1,sigma2,ib)

xfcr=zeros(indn/2,loci);
xmcr=zeros(indn/2,loci);

%calculation of beak size of the females:

for ll=1:indn/2
for l=1:loci
```

```
    if xfcr1(ll,l)==1 && xfcr2(ll,l)==1
        xfcr(ll,l)=ib;
    else if xfcr1(ll,l)==1 && xfcr2(ll,l)==0
            xfcr(ll,l)=1;
    else if xfcr1(ll,l)==0 && xfcr2(ll,l)==1
            xfcr(ll,l)=1;
    else if xfcr1(ll,l)==0 && xfcr2(ll,l)==0
            xfcr(ll,l)=0;
    end
    end
    end
    end
end
end
xf1=xdel.*xfcr;
xf=sum(xf1');

%calculation of beak size of the males:

for mm=1:indn/2
for m=1:loci
    if xmcr1(mm,m)==1 && xmcr2(mm,m)==1
        xmcr(mm,m)=ib;
    else if xmcr1(mm,m)==1 && xmcr2(mm,m)==0
            xmcr(mm,m)=1;
    else if xmcr1(mm,m)==0 && xmcr2(mm,m)==1
            xmcr(mm,m)=1;
    else if xmcr1(mm,m)==0 && xmcr2(mm,m)==0
            xmcr(mm,m)=0;
    end
    end
    end
    end
end
end
xm1=xdel.*xmcr;
xm=sum(xm1');

totx=[xf xm];

avgx=mean(totx);
varx=var(totx);

%calculation of number of individuals in a niche

p1=0;
p2=0;
for k=1:indn
    diff(k)=totx(k)-1;
    dif(k)=round(diff(k),6);
    if dif(k)<0
        p1=p1+1;
    else if dif(k)>0
            p2=p2+1;
    else if dif(k)==0
            p1=p1+0;
```

```
            p2=p2+0;
    end
    end
    end
end
nn1=p1;
nn2=p2;

intp=((sqrt(2)*meant)+(sqrt(2)*meano))/(sqrt(2)+sqrt(2));
fitintp=exp(-(intp-meano)^2/(2*sigma1*sigma1));

ff=zeros(indn/2,1);
fm=zeros(indn/2,1);

%calculation of fitness of the females
for i=1:indn/2
    xfeff(i)=round(xf(i),6);

    if xfeff(i)<1
        ff(i)=exp(-((xf(i)-meano)^2)/(2*sigma1*sigma1))*(1-
(p1/indn));
    else if xfeff(i)>1
            ff(i)=exp(-((xf(i)-meant)^2)/(2*sigma2*sigma2))*(1-
(p2/indn));
    else ff(i)=fitintp;
    end
    end
end
fitf=ff;


%calculation of fitness of the males
for i=1:indn/2
     xmeff(i)=round(xm(i),6);
    if xmeff(i)<1
        fm(i)=exp(-((xm(i)-meano)^2)/(2*sigma1*sigma1))*(1-
(p1/indn));
    else if xmeff(i)>1
            fm(i)=exp(-((xm(i)-meant)^2)/(2*sigma2*sigma2))*(1-
(p2/indn));
    else fm(i)=fitintp;
    end
    end
end
fitm=fm;

end


function[tm]=tcalc(tmcr1,tmcr2,indn,loci,tdel,it)

tmcr=zeros(indn/2,loci);

%calculation of t:
for mm=1:indn/2
for m=1:loci
```

```
    if tmcr1(mm,m)==1 && tmcr2(mm,m)==1
        tmcr(mm,m)=it;
    else if tmcr1(mm,m)==1 && tmcr2(mm,m)==0
            tmcr(mm,m)=1;
    else if tmcr1(mm,m)==0 && tmcr2(mm,m)==1
            tmcr(mm,m)=1;
    else if tmcr1(mm,m)==0 && tmcr2(mm,m)==0
            tmcr(mm,m)=0;
    end
    end
    end
    end
end
end
tm1=tdel.*tmcr;
tm=sum(tm1');


end


function[pf]=pcalc(pfcr1,pfcr2,indn,loci,pdel,ip)

pfcr=zeros(indn/2,loci);

%calculation of p:
for ll=1:indn/2
for l=1:loci
    if pfcr1(ll,l)==1 && pfcr2(ll,l)==1
        pfcr(ll,l)=ip;
    else if pfcr1(ll,l)==1 && pfcr2(ll,l)==0
            pfcr(ll,l)=1;
    else if pfcr1(ll,l)==0 && pfcr2(ll,l)==1
            pfcr(ll,l)=1;
    else if pfcr1(ll,l)==0 && pfcr2(ll,l)==0
            pfcr(ll,l)=0;
    end
    end
    end
    end
end
end
pf1=pdel.*pfcr;
pf=sum(pf1');

end


function[xfemcr1,xfemcr2,xmalcr1,xmalcr2]=xmat(xfcr1,xfcr2,xmcr1,xmc
r2,indn,loci,fmat,mmat);

%first half of the individuals generated are females, second half
males%

%genotype of the chromosome from the female:
for i=1:indn
```

```
    f=fmat(i); %f is the female that mates in the ith mating event.
    for j=1:loci
        r1=rand;
        r2=rand;
        if r1<0.5 && r2>0.00001
            crf1(i,j)=xfcr1(f,j);
        else if r1<0.5 && r2<0.00001
                crf1(i,j)=xfcr2(f,j);
        else if r1>0.5 && r2>0.00001
                crf1(i,j)=xfcr2(f,j);
        else crf1(i,j)=xfcr1(f,j);
        end
        end
    end
    end
end
xfemcr1=crf1(1:indn/2,:);
xmalcr1=crf1(indn/2+1:indn,:);

%genotype of the chromosome from the male:
for i=1:indn
    m=mmat(i); %m is the male that mates in the ith mating event%
    for j=1:loci
        r3=rand;
        r4=rand;
        if r3<0.5 && r4>0.00001
            crf2(i,j)=xmcr1(m,j);
        else if r3<0.5 && r4<0.00001
                crf2(i,j)=xmcr2(m,j);
        else if r3>0.5 && r4>0.00001
                crf2(i,j)=xmcr2(m,j);
        else crf2(i,j)=xmcr1(m,j);
        end
        end
    end
    end
end
xfemcr2=crf2(1:indn/2,:);
xmalcr2=crf2(indn/2+1:indn,:);

end


function[tfemcr1,tfemcr2,tmalcr1,tmalcr2]=tmat(tfcr1,tfcr2,tmcr1,tmc
r2,indn,loci,fmat,mmat);

%first half of the individuals generated are females, second half
males%

%genotype of the chromosome from the female:
for i=1:indn
    f=fmat(i);
    for j=1:loci
        r1=rand;
        r2=rand;
        if r1<0.5 && r2>0.00001
```

```
            crf1(i,j)=tfcr1(f,j);
        else if r1<0.5 && r2<0.00001
                crf1(i,j)=tfcr2(f,j);
        else if r1>0.5 && r2>0.00001
                crf1(i,j)=tfcr2(f,j);
        else crf1(i,j)=tfcr1(f,j);
        end
        end
    end
    end
end
tfemcr1=crf1(1:indn/2,:);
tmalcr1=crf1(indn/2+1:indn,:);

%genotype of the chromosome from the male:
for i=1:indn
    m=mmat(i);
    for j=1:loci
        r3=rand;
        r4=rand;
        if r3<0.5 && r4>0.00001
            crf2(i,j)=tmcr1(m,j);
        else if r3<0.5 && r4<0.00001
                crf2(i,j)=tmcr2(m,j);
        else if r3>0.5 && r4>0.00001
                crf2(i,j)=tmcr2(m,j);
        else crf2(i,j)=tmcr1(m,j);
        end
        end
    end
    end
end
tfemcr2=crf2(1:indn/2,:);
tmalcr2=crf2(indn/2+1:indn,:);

end


function[pfemcr1,pfemcr2,pmalcr1,pmalcr2]=pmat(pfcr1,pfcr2,pmcr1,pmc
r2,indn,loci,fmat,mmat);

%first half of the individuals generated are females, second half
males%

%genotype of the chromosome from the female:
for i=1:indn
    f=fmat(i);
    for j=1:loci
        r1=rand;
        r2=rand;
        if r1<0.5 && r2>0.00001
            crf1(i,j)=pfcr1(f,j);
        else if r1<0.5 && r2<0.00001
                crf1(i,j)=pfcr2(f,j);
        else if r1>0.5 && r2>0.00001
                crf1(i,j)=pfcr2(f,j);
```

```
        else crf1(i,j)=pfcr1(f,j);
        end
        end
    end
    end
end
pfemcr1=crf1(1:indn/2,:);
pmalcr1=crf1(indn/2+1:indn,:);

%genotype of the chromosome from the male:
for i=1:indn
    m=mmat(i);
    for j=1:loci
        r3=rand;
        r4=rand;
        if r3<0.5 && r4>0.00001
            crf2(i,j)=pmcr1(m,j);
        else if r3<0.5 && r4<0.00001
                crf2(i,j)=pmcr2(m,j);
        else if r3>0.5 && r4>0.00001
                crf2(i,j)=pmcr2(m,j);
        else crf2(i,j)=pmcr1(m,j);
        end
        end
    end
    end
end
pfemcr2=crf2(1:indn/2,:);
pmalcr2=crf2(indn/2+1:indn,:);

end
```

**Assortative mating**

```
mainfile

clc
close all
clear all

ngen=50; %number of generations
avg_asso=zeros(100,50);
dev_asso=zeros(100,50);

sig1=0.1; %width of resource distribution

for i=1:length(sig1)
   sig=sig1(i);
   ij=1;
while ij<=100
    [split(ij),avg_asso(ij,:),dev_asso(ij,:)]=symsp_asso(ngen,sig);
    ij=ij+1;
end
end


function[diff,p_asso,p_asso_dev]=symsp_asso(ngenerations,dev)

nloci=20;  % number of loci that control x,t,p and assortativeness%
nind=1000;  %number of individuals%

alpha=5; %strength of sexual selection
delx=2/(2*nloci); %contribution of each locus towards beak size
delt=0.8/(2*nloci); %contribution of each locus towards investment strategy of
males
delp=0.4/(2*nloci); %contribution of each locus towards choosiness of females
dela=1/(2*nloci); %contribution of each locus towards assortativeness
sump=0.4; %maximum permissible choosiness
%width of resouce distribution
sig1=dev;
sig2=dev;

matm=zeros(nind,1); %array of males that mate%
matf=zeros(nind,1); %array of females that mate%

xtrack=zeros(ngenerations,nind); %beak size of individuals of every generation%
ttrack=zeros(ngenerations,nind/2); %investment strategies of the males of every
generation%
ptrack=zeros(ngenerations,nind/2); %choosiness of the females of every generation%
mtrack=zeros(ngenerations,nind);
ftrack=zeros(ngenerations,nind);

fpcr1=zeros(nind/2,nloci); %first strand of female's gene - p
fpcr2=zeros(nind/2,nloci); %second strand of female's gene - p
mpcr1=zeros(nind/2,nloci); %first strand of male's gene - p
mpcr2=ones(nind/2,nloci); %second strand of male's gene - p

ftcr1=zeros(nind/2,nloci); %first strand of female's gene - t
ftcr2=ones(nind/2,nloci); %second strand of female's gene - t
mtcr1=zeros(nind/2,nloci); %first strand of male's gene - t
mtcr2=zeros(nind/2,nloci); %second strand of male's gene - t
```

```
fxcr1=zeros(nind/2,nloci); %first strand of female's gene - x
fxcr2=ones(nind/2,nloci); %second strand of female's gene - x
mxcr1=zeros(nind/2,nloci); %first strand of male's gene - x
mxcr2=ones(nind/2,nloci); %second strand of male's gene - x

fasscr1=zeros(nind/2,nloci); %first strand of female's gene - assortativess
fasscr2=zeros(nind/2,nloci); %second strand of female's gene - assortativeness
masscr1=zeros(nind/2,nloci); %first strand of male's gene - assortativeness
masscr2=ones(nind/2,nloci); %second strand of male's gene - assortativeness


%creation of normal distributions that indicate disruptive selection%
mean1=0.5;
mean2=1.5;


for i=1:ngenerations

    msmallbs=0; %number of individuals with beak size < 1
    mbigbs=0; %number of individuals with beak size > 1
    ngen(i)=i;

    %calculation of assortativeness of the females
    [fscore,assf]=assocalc(fasscr1,fasscr2,masscr1,masscr2,nind,dela);
    p_asso(i)=mean(fscore);
    p1_ass(i)=mean(assf);
    p_asso_dev(i)=std(fscore);


[ff,fm,xavg,xvar,xtot,np1,np2,fx,mx]=fcalc_ass(fxcr1,fxcr2,nind,mxcr1,mxcr2,nloci,
delx,mean1,mean2,sig1,sig2); %function that calculates beak size and fitness of
the individuals


    avgxx(i)=xavg;
    varxx(i)=xvar;
    fitness=cat(1,ff,fm);
    avgf(i)=mean(fitness);

    femfit=ff;
    recif=femfit.^(-1);
    malfit=fm;

    [fp]=pcalc_ass(fpcr1,fpcr2,nind,nloci,delp); %function that calculates p of
every female%
    avgp(i)=mean(fp);
    % calculation of probability that a female finds a partner
    fcost=(exp(-8*fp.*fp.*fp/(sump)^3)).*femfit';
    sumfcost=sum(fcost);
    fprob1=fcost/sumfcost;

    sumf=0;
    for x=1:nind/2
        sumf=sumf+fprob1(x);
        fprob(x)=sumf;
    end
```

```
    [mt]=tcalc_ass(mtcr1,mtcr2,nind,nloci,delt); %function that calculates t of
every male%
    avgt(i)=mean(mt);

    % calculation of probability that a male escapes predation%
    for jj=1:nind/2
        predprop(jj)=exp(-mt(jj)/(1-mt(jj)));
    end

    summcost=sum(predprop);
    predprob=predprop/summcost;
    summ=0;
    for y=1:nind/2
        summ=summ+predprob(y);
        mprob(y)=summ;
    end

    if i>=1
        xtrack(i,:)=xtot;
        ttrack(i,:)=mt;
        ptrack(i,:)=fp;
    end

    nmatef=0.000*ones(nind/2,1);

    %meeting and mating events%
    for p=1:nind
        r1=rand; %selection of female that mates%
        ii=1;
        while r1>fprob(ii)
            ii=ii+1;
        end
        matf(p)=ii;
        nmatef(ii)=nmatef(ii)+1;

     if fscore(ii)>0
         prb=fscore(ii); %assortativness of the female that is selected for mating
         bsc=fx(ii); %beak size of the female selected for mating
        if bsc<1
        for q=1:nind/2
            if mx(q)<1
            fdep(q)=prb*(1-mt(q))*malfit(q)*mprob(q)/(1-prb); %probability that
the selected female meets males whose beak sizes are similar to its own
            matingp(q)=exp(mt(q)*malfit(q)*fp(ii)*alpha); %probability that the
selected female meets mates with males whose beak sizes are similar to its own
            else
            fdep(q)=(1-prb)*(1-mt(q))*malfit(q)*mprob(q)/prb; %probability that
the selected female meets males whose beak sizes are dissimilar to its own
            matingp(q)=exp(mt(q)*malfit(q)*fp(ii)*alpha); %probability that the
selected female mates with males whose beak sizes are dissimilar to its own
            end
        end
        else if bsc>1
            for q=1:nind/2
            if mx(q)>1
            fdep(q)=prb*(1-mt(q))*malfit(q)*mprob(q)/(1-prb); %probability that
the selected female meets males whose beak sizes are similar to its own
            matingp(q)=exp(mt(q)*malfit(q)*fp(ii)*alpha); %probability that the
selected female meets mates with males whose beak sizes are similar to its own
```

```
            else
            fdep(q)=(1-prb)*(1-mt(q))*malfit(q)*mprob(q)/prb; %probability that
the selected female meets males whose beak sizes are dissimilar to its own
            matingp(q)=exp(mt(q)*malfit(q)*fp(ii)*alpha); %probability that the
selected female mates with males whose beak sizes are dissimilar to its own
            end
            end
        end
        end
     else
         for q=1:nind/2
            fdep(q)=0.0001*(1-mt(q))*malfit(q)*mprob(q); %product of the selected
female's fitness with all the males' fitness"
            matingp(q)=exp(mt(q)*malfit(q)*fp(ii)*alpha); %mating probability of
the selected female with all the males in the population
         end
     end

            sumfdep=sum(fdep);
            fdepp=fdep/sumfdep;
            summatp=sum(matingp);
            probmating=matingp/summatp;

            sumffdep=0;
            summating=0;

            for pp=1:nind/2
              sumffdep=sumffdep+fdepp(pp);
              fitnessprob(pp)=sumffdep;
              summating=summating+probmating(pp);
              matprb(pp)=summating;
            end

        %identification of meeting event, and then the mating event%
        iii=1;
        iiii=1;
        while iiii==1
            r2=rand;
            r3=rand;
         while r2>fitnessprob(iii)
            iii=iii+1;
         end
         ev=iii;
         if r3>matprb(ev)
            iiii=1;
         else
             iiii=0;

         end
        end
        matm(p)=ev;
    end

    for b=1:nind
        indf=matf(b);
        indm=matm(b);
        matfemx(b)=fx(indf);
        matmalx(b)=mx(indm);
    end
```

```
    if i>=1
        mtrack(i,:)=matmalx;
        ftrack(i,:)=matfemx;
    end



    %generation of the genotypes of offspring of next generation

[fempcr1,fempcr2,malpcr1,malpcr2]=pmat_ass(fpcr1,fpcr2,mpcr1,mpcr2,nind,nloci,matf
,matm);

[femtcr1,femtcr2,maltcr1,maltcr2]=tmat_ass(ftcr1,ftcr2,mtcr1,mtcr2,nind,nloci,matf
,matm);

[femxcr1,femxcr2,malxcr1,malxcr2]=xmat_ass(fxcr1,fxcr2,mxcr1,mxcr2,nind,nloci,matf
,matm);

[asscr1_f,asscr2_f,asscr1_m,asscr2_m]=assmat(fasscr1,fasscr2,masscr1,masscr2,nind,
nloci,matf,matm);

    %reassignment of female and male genotypes%
    fpcr1=fempcr1;
    fpcr2=fempcr2;
    mpcr1=malpcr1;
    mpcr2=malpcr2;

    ftcr1=femtcr1;
    ftcr2=femtcr2;
    mtcr1=maltcr1;
    mtcr2=maltcr2;

    fxcr1=femxcr1;
    fxcr2=femxcr2;
    mxcr1=malxcr1;
    mxcr2=malxcr2;

    fasscr1=asscr1_f;
    fasscr2=asscr2_f;
    masscr1=asscr1_m;
    masscr2=asscr2_m;
end

femxf=ftrack(ngenerations-1,:);
malxf=mtrack(ngenerations-1,:);
matxf=[femxf malxf];

%calculation of intensity of split
diff=1;
bs1=0;
for pp=1:(2*nind)
    if matxf(pp)==1
        bs1=bs1+1;
    else
        bs1=bs1+0;
    end
end
```

```
if bs1~=0
    diff=0;
else
nsmallx=0;
nbigx=0;
 for hh=1:(nind*2)
    if matxf(hh)<1
        nsmallx=nsmallx+1;
    else if matxf(hh)>1
            nbigx=nbigx+1;
    end
    end
 end

  if nsmallx==0 && nbigx~=0
    diff=-1;
  else if nsmallx~=0 && nbigx==0
          diff=-1;
  else if nsmallx<2 || nbigx<2
          diff=-1;
  else if (nsmallx~=0 && nbigx~=0) && (nsmallx>2 && nbigx>2)

      smallx=zeros(nsmallx,1);
      bigx=zeros(nbigx,1);

      fmatx=sort(matxf);

      smallx=(fmatx(:,1:nsmallx))';
      bigx=(fmatx(:,(nsmallx+1):end))';

      pd1=fitdist(smallx,'Normal');
      m1=mean(pd1);
      dev1=std(pd1);
      pd2=fitdist(bigx,'Normal');
      m2=mean(pd2);
      dev2=std(pd2);

      diff=(m2-m1)/(dev2+dev1);
  end
  end
  end
  end
end
end

function[totalf,scoref]=assocalc(cr1assf,cr2assf,cr1assm,cr2assm,indn,adel)

scoref=0;
scorem=0;
femasscr=cr1assf+cr2assf;
malasscr=cr2assm+cr1assm;

totalf=adel*sum(femasscr'); %assortativeness of the females


for i=1:indn/2
    if totalf(i)>0
        scoref=scoref+1;
```

```
    end
end

end
```
