## Supplementary Figures for "When is sympatric speciation a possible evolutionary outcome?"

### Supplementary Material

**Effect of increase in DS and SS on split intensity, beak size, choosiness and investment strategy.**

#### A. Intensity of split

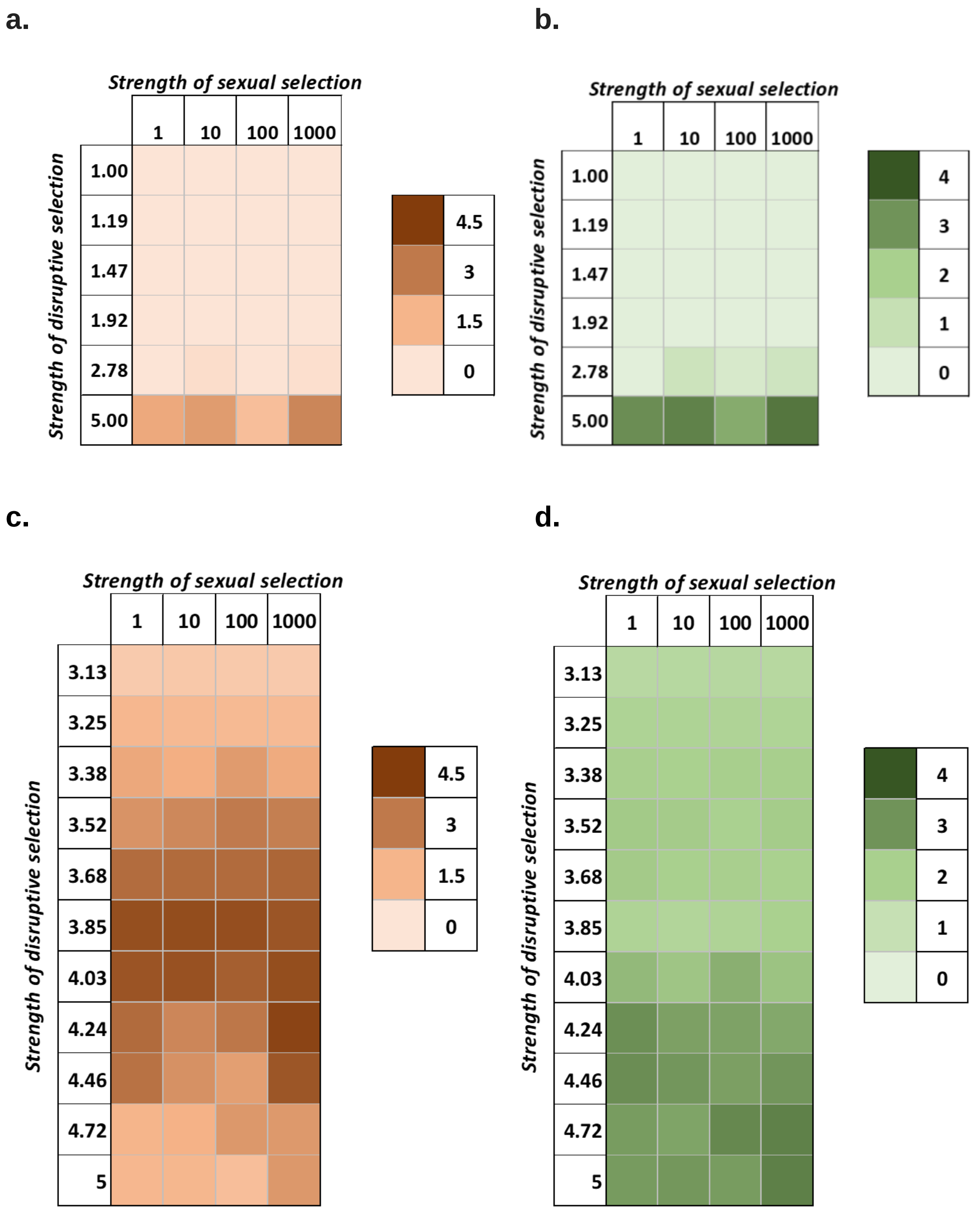


**Figure S1. Mean intensity of split (a and c) and standard deviation (b and d) (calculated as average obtained considering all the evolutionary outcomes) at the end of 50 generations.**

#### B. Beak size

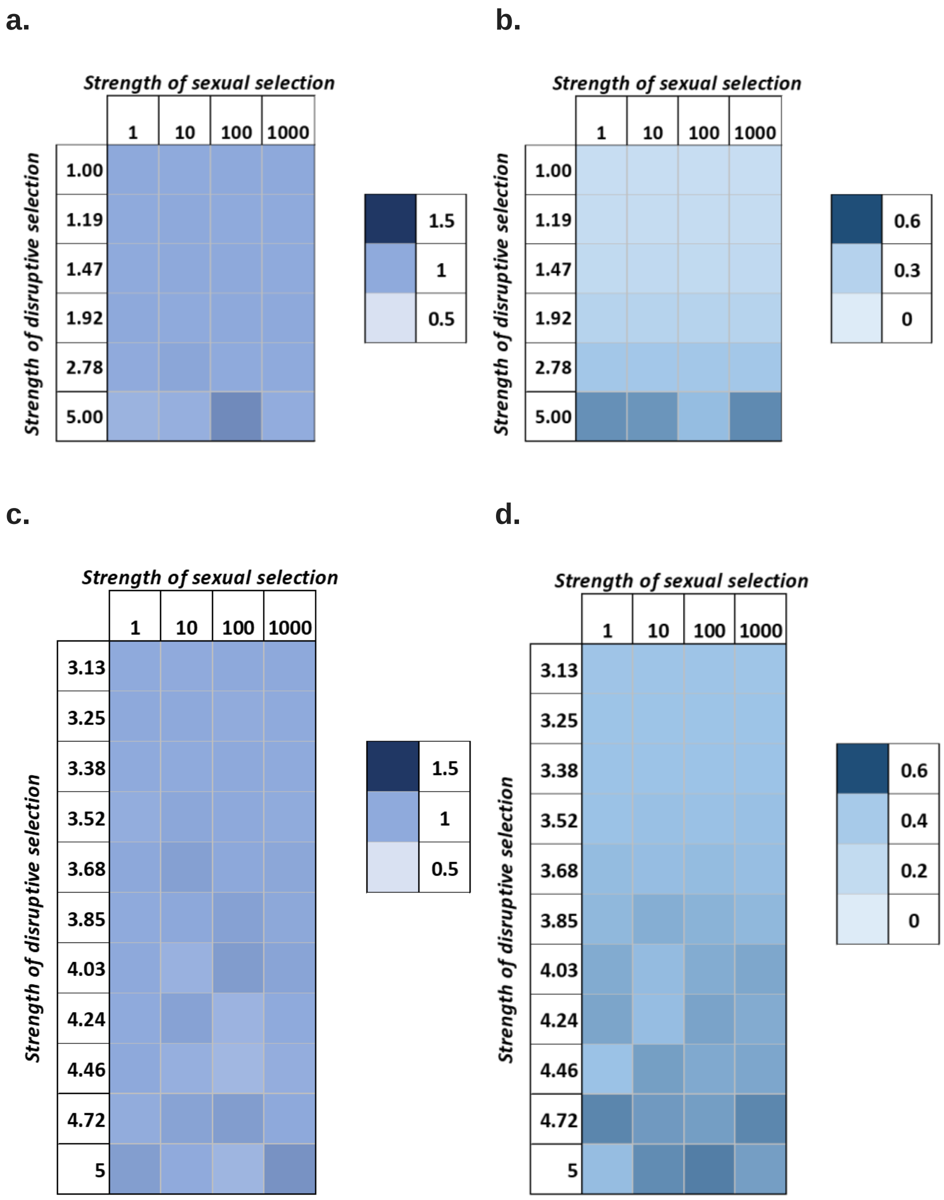


**Figure S2: Mean beak sizes (a and c) and standard deviations (b and d) of the population at the end of 50 generations.**

#### C. Choosiness

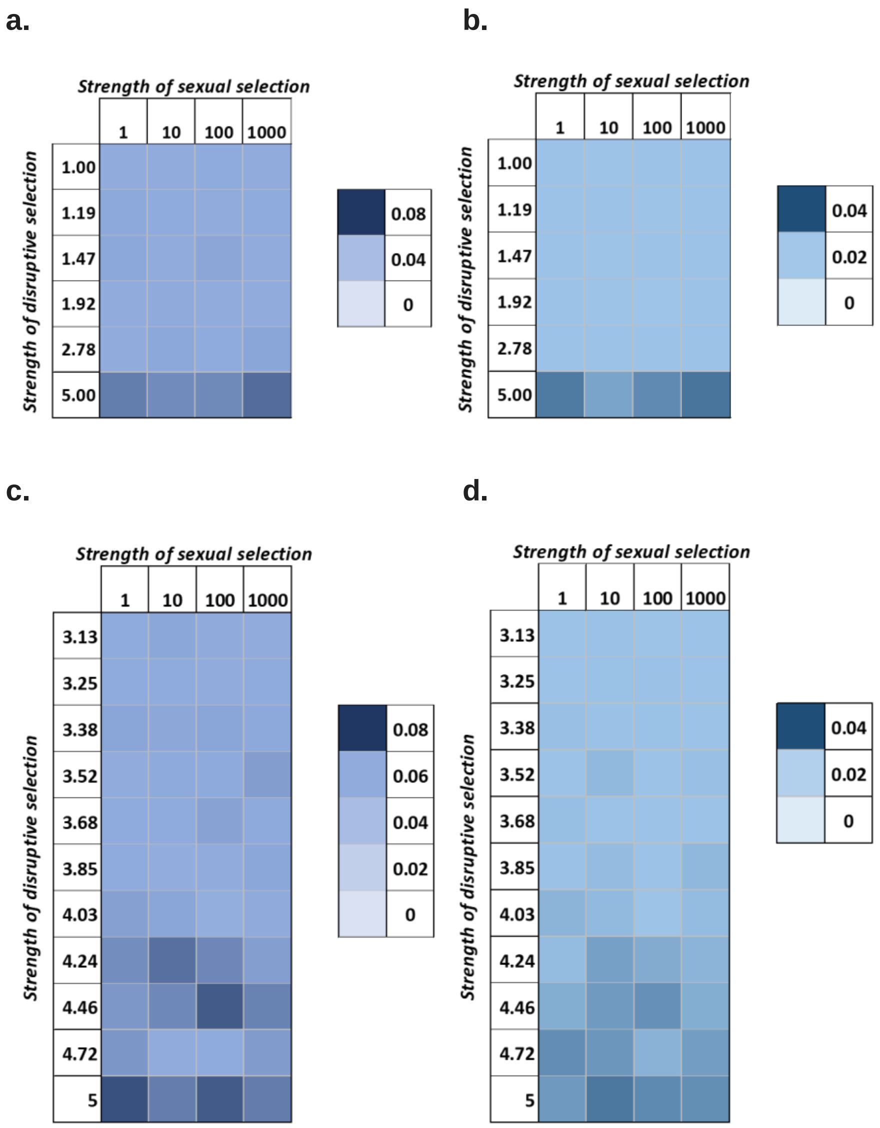


**Figure S3: Mean choosiness (a and c) and standard deviations (b and d) of the population at the end of 50 generations.**

#### D. Investment strategy

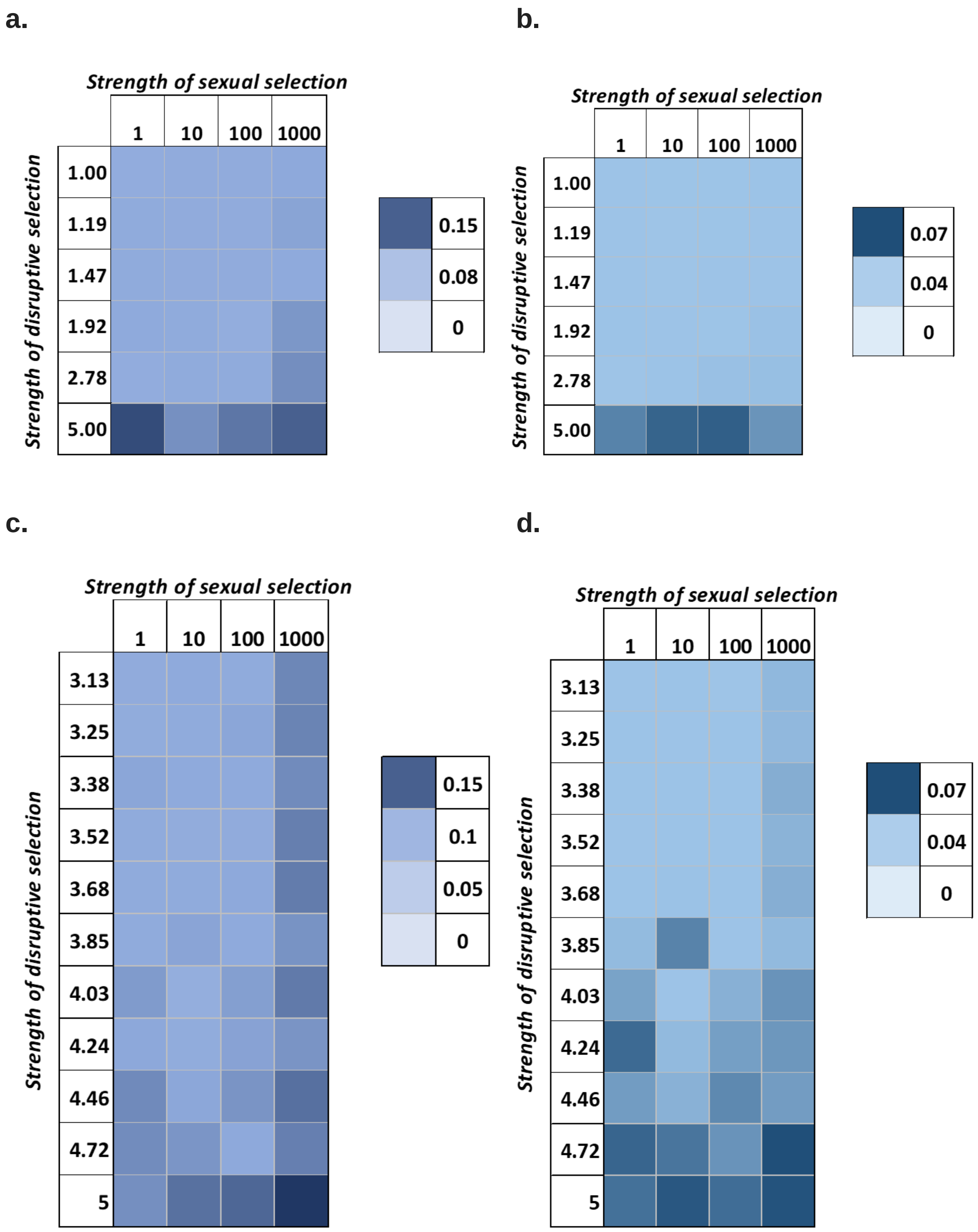


**Figure S4: Mean investment strategy (a and c) and standard deviations (b and d) of the population at the end of 50 generations.**

**Effect of altering genetic architecture on the intensity of split and beak size**

#### 1. Null case

##### A. Intensity of split

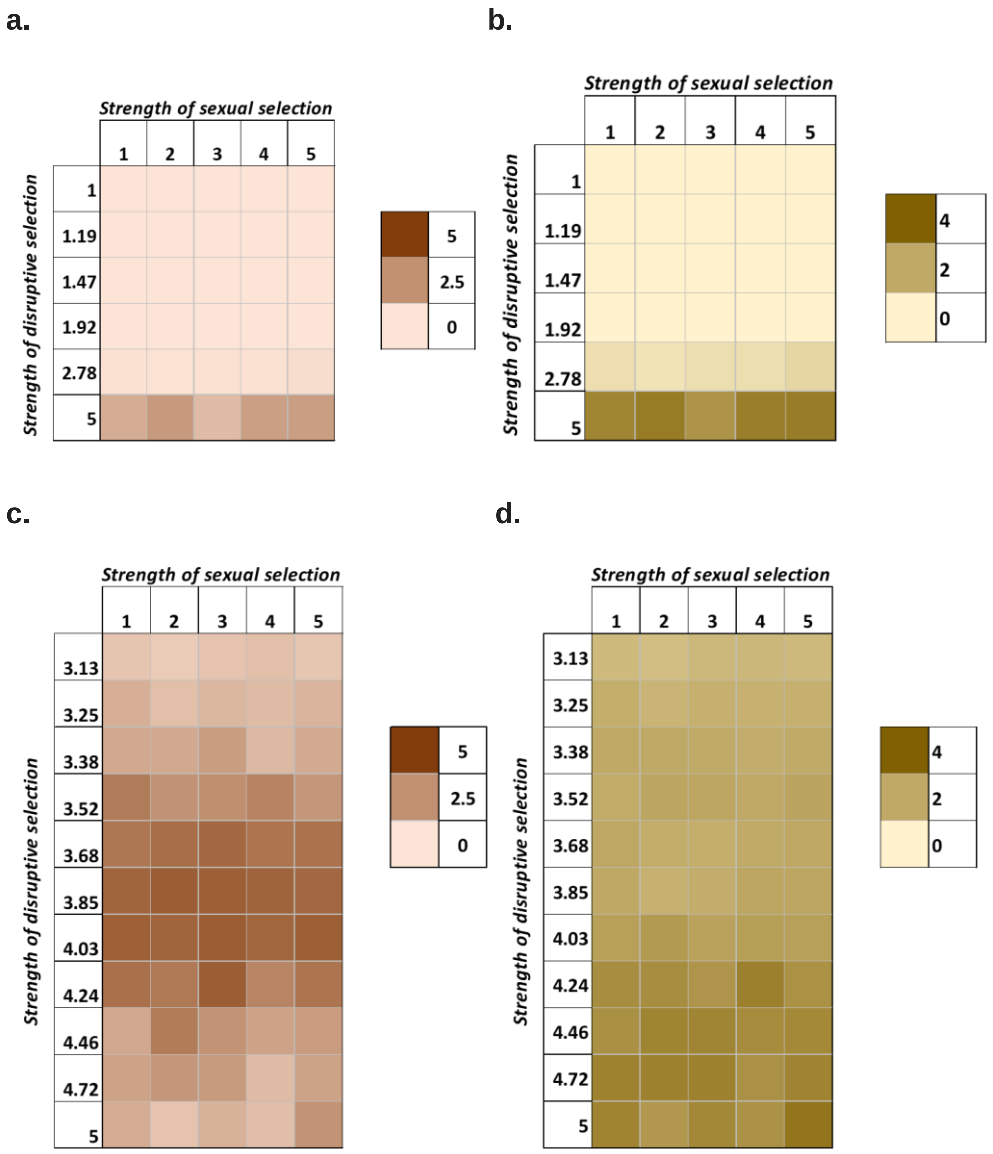


**Figure S5. Mean intensity of split (a and c) and standard deviation (b and d) (calculated as average obtained considering all the evolutionary outcomes) at the end of 50 generations.**

#### B. Beak size

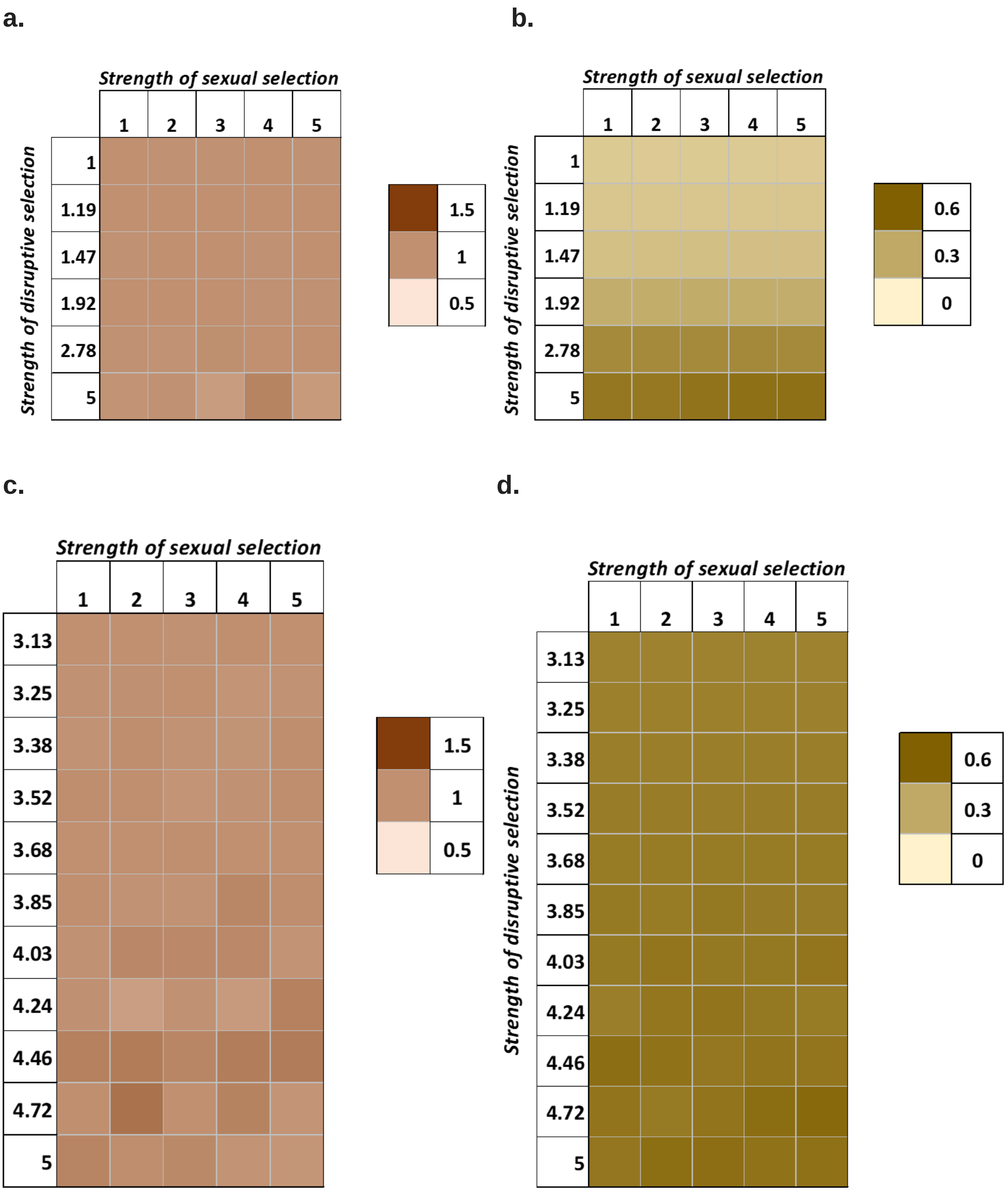


**Figure S6: Mean beak sizes (a and c) and standard deviations (b and d) of the population at the end of 50 generations.**

#### 2. Dominance in the loci controlling beak size

##### A. Intensity of split

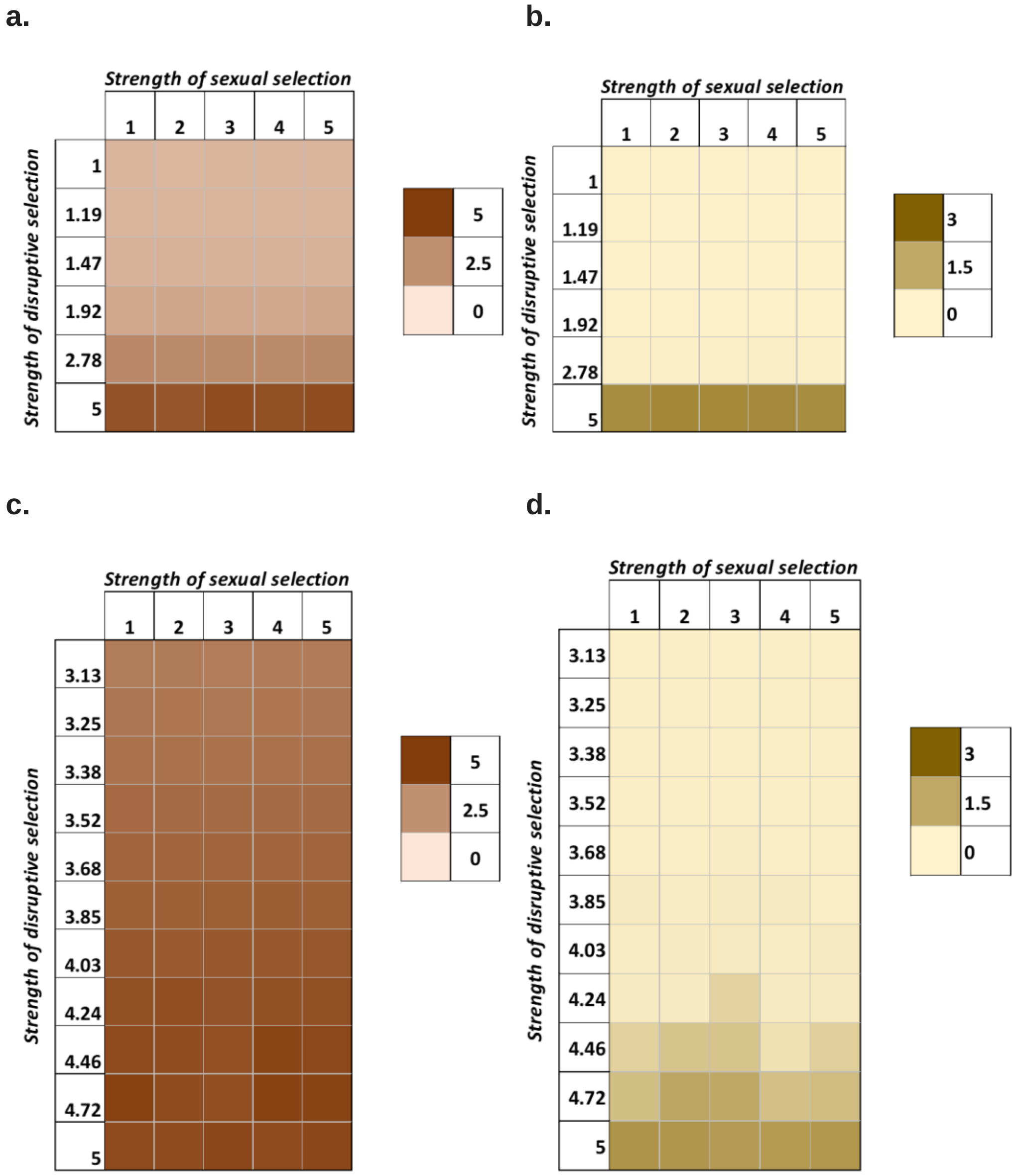


**Figure S7. Mean intensity of split (a and c) and standard deviation (b and d) (calculated as average obtained considering all the evolutionary outcomes) at the end of 50 generations.**

#### B. Beak size

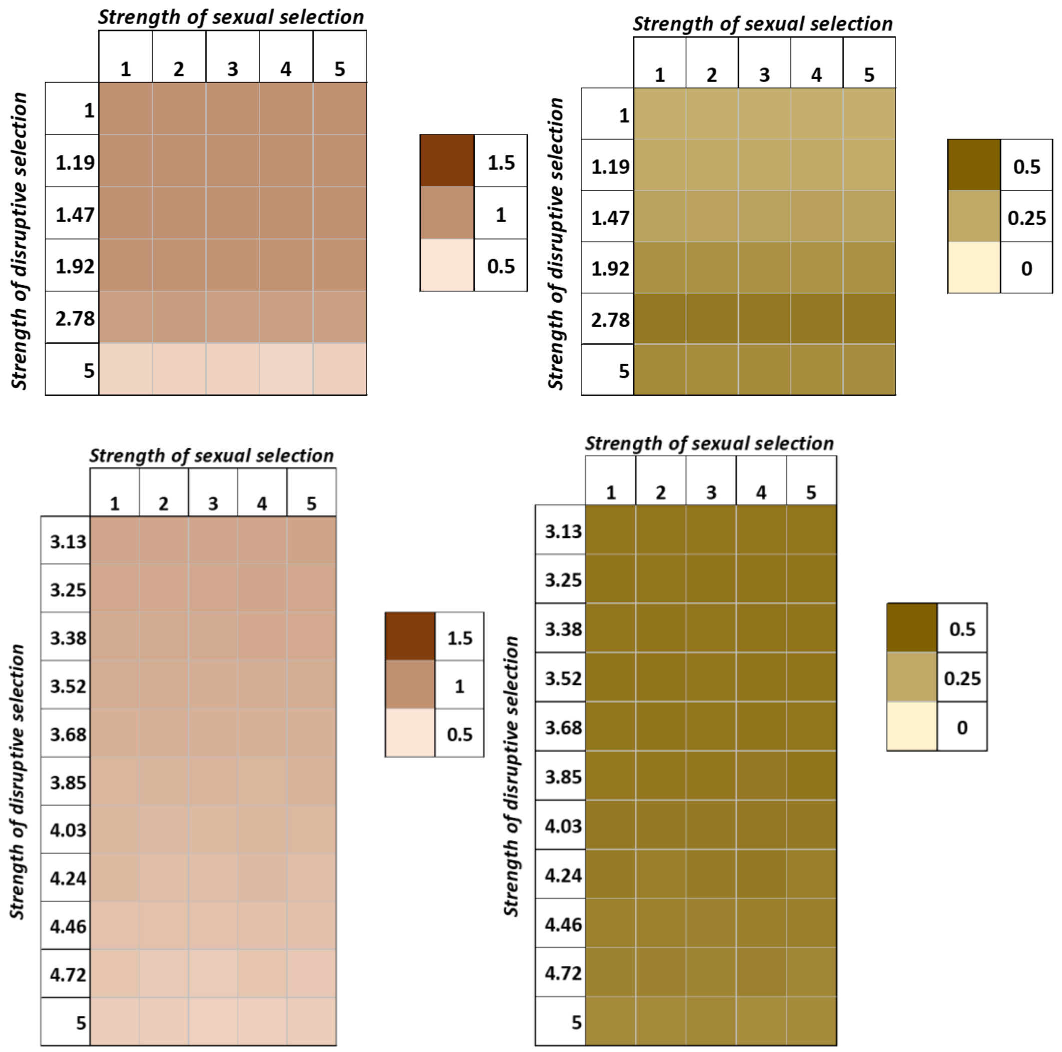


**Figure S8: Mean beak sizes (a and c) and standard deviations (b and d) of the population at the end of 50 generations.**

#### 3. Unequal contribution of loci controlling beak size

##### A. Split

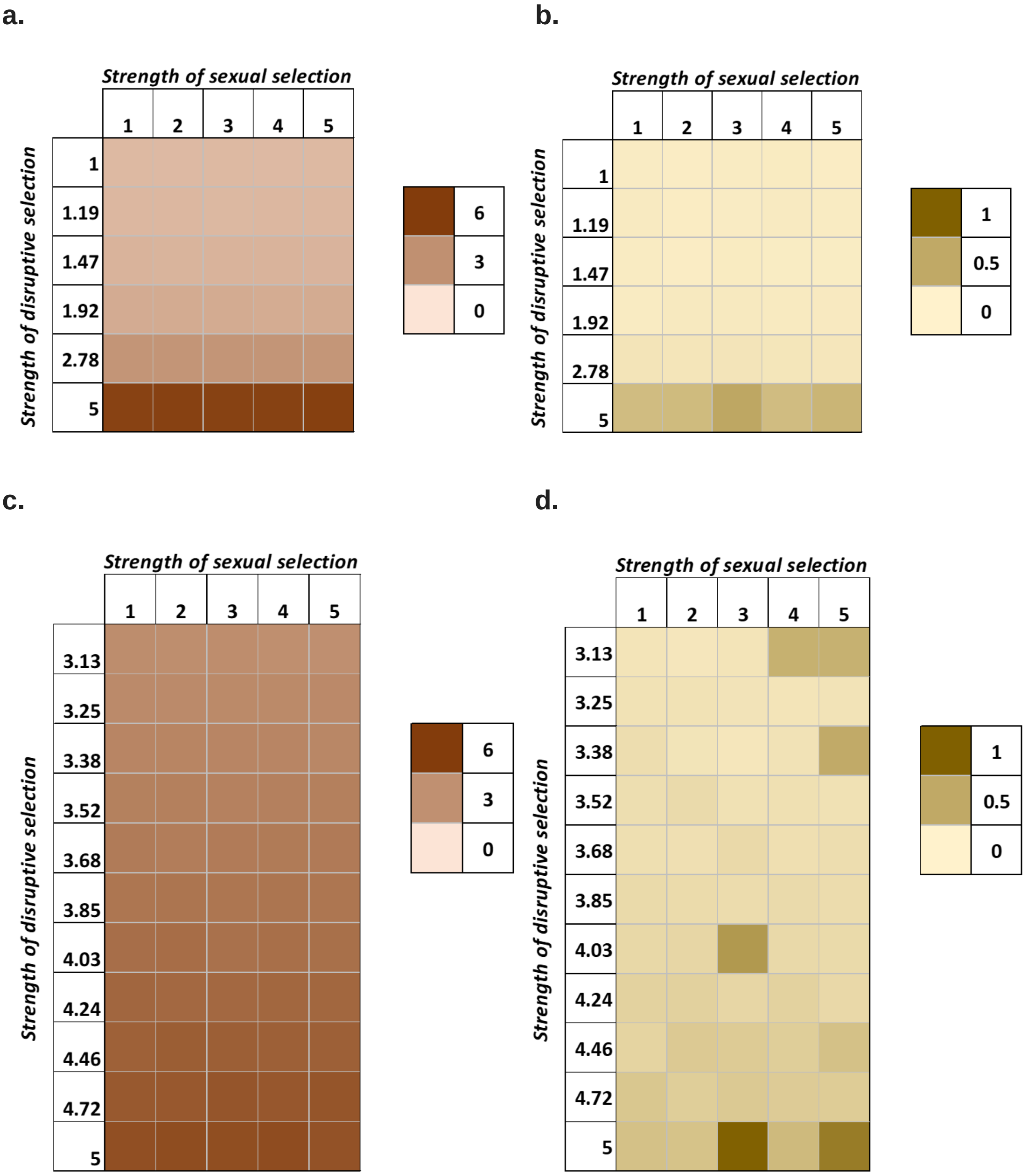


**Figure S9. Mean intensity of split (a and c) and standard deviation (b and d) (calculated as average obtained considering all the evolutionary outcomes) at the end of 50 generations.**

#### B. Beak size

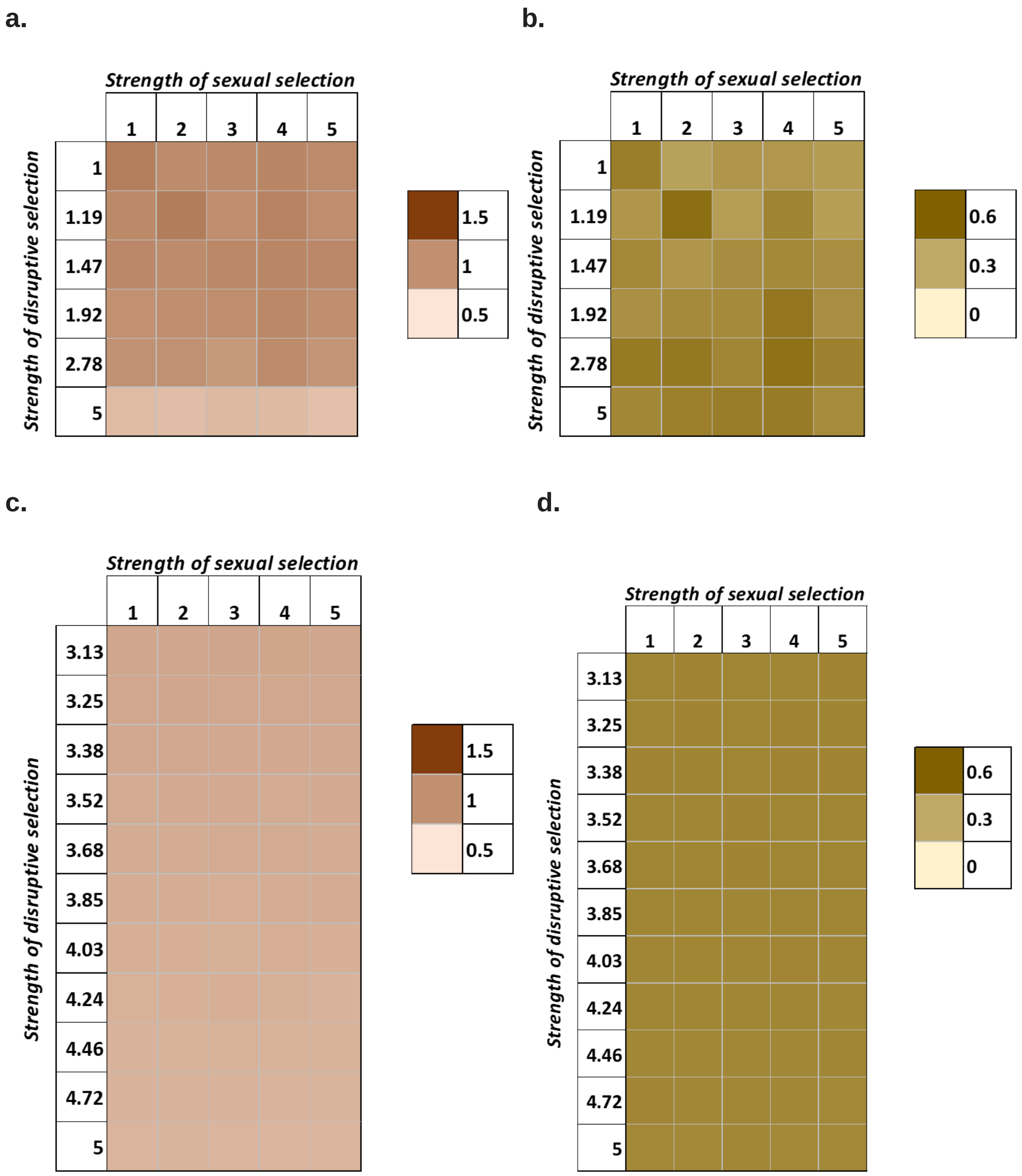


**Figure S10: Mean beak sizes (a and c) and standard deviations (b and d) of the population at the end of 50 generations.**

#### 4. Unequally contribution loci of beak size that also show dominance relationships

##### A. Split

a.

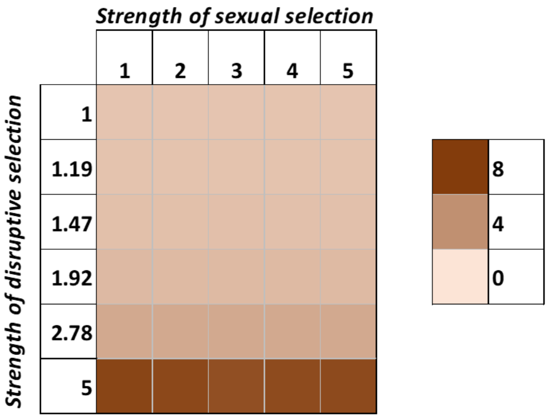


b.

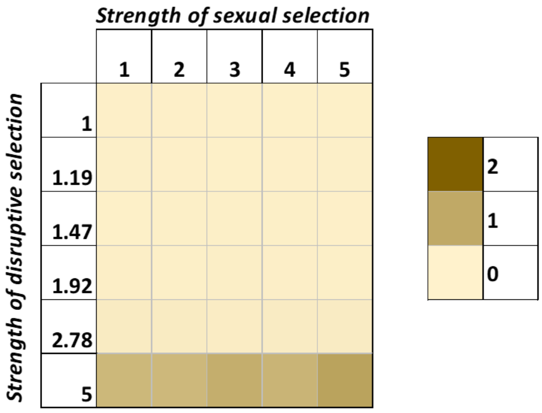


c.

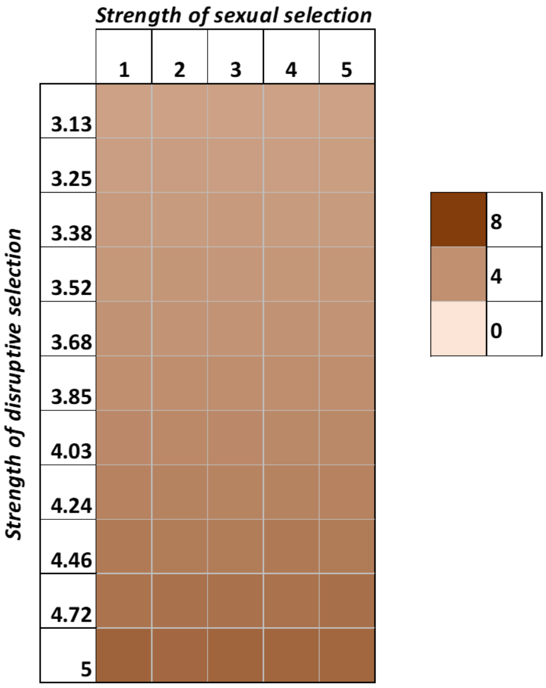


d.

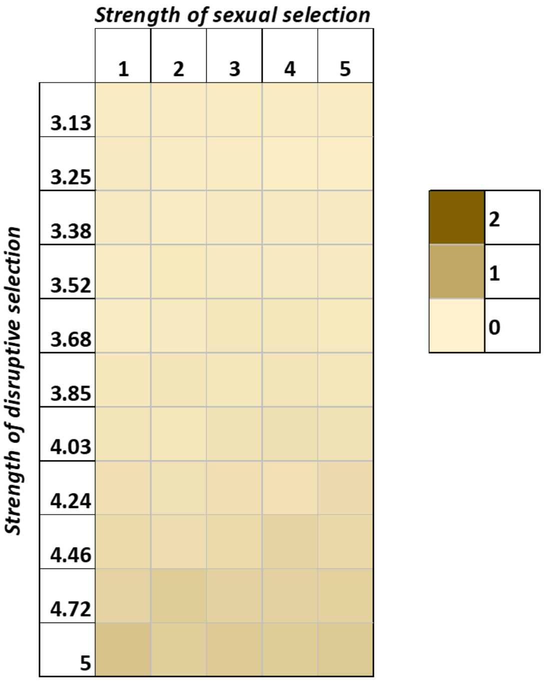


**Figure S11. Mean intensity of split (a and c) and standard deviation (b and d) (calculated as average obtained considering all the evolutionary outcomes) at the end of 50 generations.**

#### B. Beak size

a.

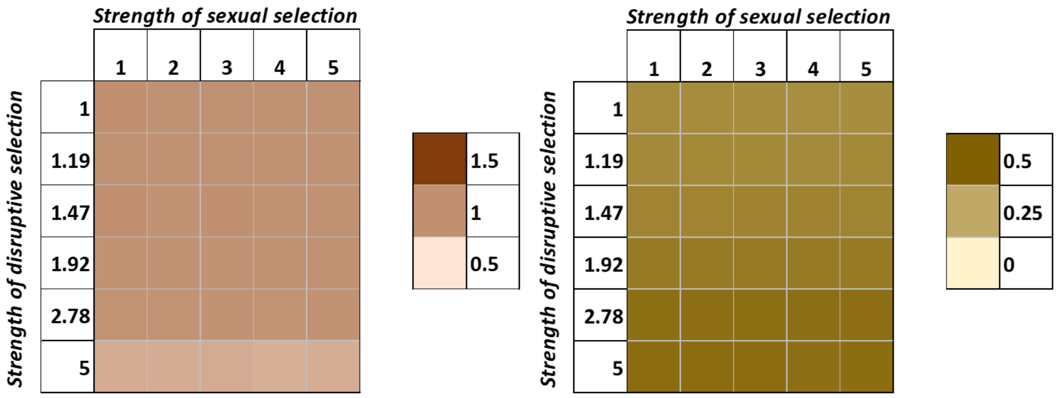


c.

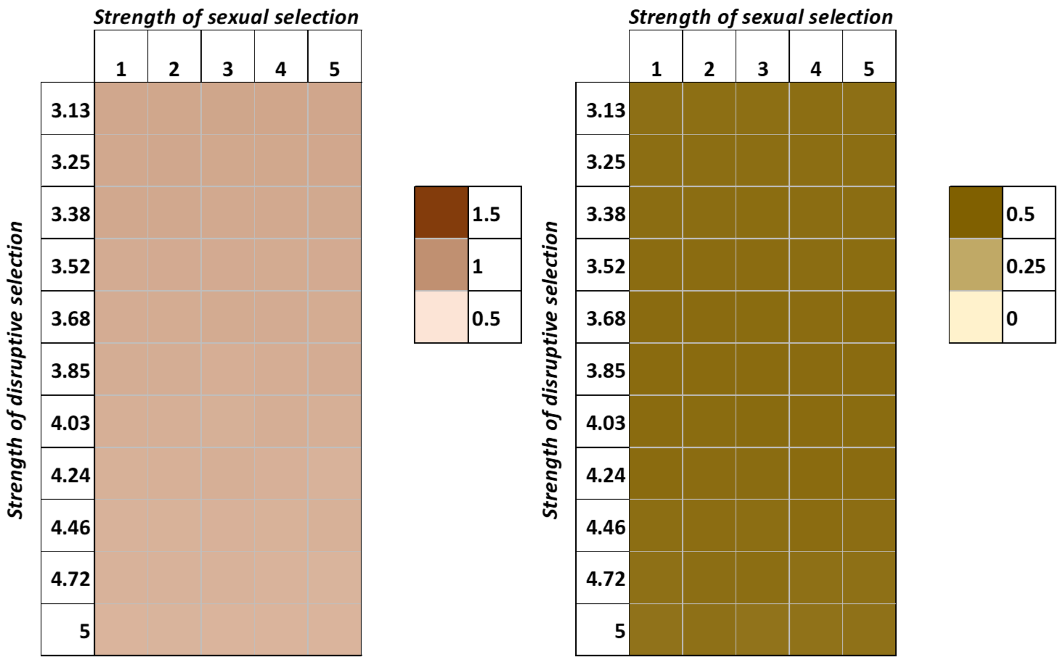


**Figure S12: Mean beak sizes (a and c) and standard deviations (b and d) of the population at the end of 50 generations.**
